# Preserved striatal innervation and motor function despite severe loss of nigral dopamine neurons following mitochondrial dysfunction induced by mtDNA mutations

**DOI:** 10.1101/2023.07.07.547791

**Authors:** Thomas Paß, Konrad M. Ricke, Pierre Hofmann, Roy Chowdhury, Yu Nie, Patrick Chinnery, Heike Endepols, Bernd Neumaier, André Carvalho, Lionel Rigoux, Sophie Steculorum, Julien Prudent, Trine Riemer, Markus Aswendt, Bent Brachvogel, Rudolf J. Wiesner

**Author notes:** Brain and Mind Research Institute, Department of Cellular and Molecular Medicine, University of Ottawa, Ottawa, Ontario, Canada (Konrad M. Ricke).

## Abstract

Degeneration of dopamine neurons in the substantia nigra and their striatal axon terminals causes cardinal motor symptoms of Parkinson’s disease (PD). In idiopathic cases, high levels of mitochondrial DNA (mtDNA) mutations associated with mitochondrial dysfunction are a central feature of these vulnerable neurons. Here we present a mouse model expressing the K320E-variant of the mitochondrial helicase Twinkle in dopamine neurons, leading to accelerated mtDNA ageing. K320E-Twinkle^DaN^ mice showed normal motor function at 20 months of age, although already ∼70% of nigral dopamine neurons had perished. The remaining neuron population still preserved ∼75% of axon terminals in the dorsal striatum, which enabled normal dopamine release. Transcriptome analysis and viral tracing confirmed compensatory axonal sprouting of surviving nigral dopamine neurons. We conclude that a small population of substantia nigra neurons can adapt to mtDNA mutations and maintain motor control in mice, holding chances for new treatment strategies in PD patients.

## Introduction

Parkinson’s disease (PD) is the most common neurodegenerative motor disease and affects millions of people worldwide^1^. Furthermore, the incidence of idiopathic cases rises with the demographic increase of life expectancy. Diagnosis of patients only occurs with the onset of cardinal motor symptoms, which are caused by the lack of striatal dopamine. At this point, already ∼30-50% of dopamine-producing neurons in the *substantia nigra* (SN)^1–3^ and ∼50-70% of their axon projections in the striatum are lost^4–7^.

Other dopaminergic neuron populations, however, are spared^8–10^. This selective vulnerability is linked to cell type-specific factors: SN dopamine neurons possess extraordinary long, extensively branched and unmyelinated axons, which form an enormous number of synapses^11^ and have thus high energetic demands^12^. In addition, pacemaker activity of SN dopamine neurons generates oscillatory increases in free cytosolic Ca^2+^ levels, which are linked to cell death in the presence of mitochondrial dysfunction^13, 14^, especially disturbing mitochondrial redox homeostasis^15^.

To explain the ageing-related increase in the frequency of developing idiopathic PD, two major hypotheses were put forward: the oligomerization and aggregation of α-synuclein, leading to Lewy body formation, and mitochondrial dysfunction. α-synuclein pathology may result from its modest, long-term upregulation in idiopathic PD^16, 17^, however the exact causes for its elevation in non-familial PD remain elusive. On the other hand, mitochondrial respiratory chain deficiency is a well-known age-related feature of SN dopamine neurons in PD patients^18^. This defect is caused by the accumulation of mitochondrial DNA (mtDNA) deletions and gene duplications (indels) over time. Using the newly developed PCR method, two groups showed 30 years ago that such indels accumulate during ageing, and especially in brain regions enriched in dopamine neurons^19, 20^. More convincingly, analysis of single SN dopamine neurons showed that ageing leads to the accumulation of mtDNA deletions^21^ and even more severely in PD^22^. We have shown *in vivo* and *in vitro* that dopamine metabolism is highly mutagenic for mtDNA^23, 24^. Recently, the group of C. Tzoulis found that the total copy number of mtDNA is increased in dopamine neurons of healthy individuals during normal ageing, thus compensating the effect of defective molecules. In PD patients, on the other hand, this compensatory response did not occur^25^.

Mitochondrial complex I inhibitors like MPTP and Rotenone or genetic models of mitochondrial dysfunction have been widely used to model the disease. However, these approaches usually lead to rapid neuron death and do not reflect the slowly progressing pathology in PD, including the ageing-associated accumulation of mtDNA deletions. We have developed a mouse model overexpressing a dominant-negative mutant (K320E) of the single mitochondrial helicase Twinkle in various cell types of mice (Rosa26-STOP-loxP-Cre System). Expression of K320E-Twinkle accelerates the accumulation of mtDNA indels in non-dividing cell types, such as cardiomyocytes^26^ and skeletal muscle cells^27^, while it leads to a loss of mtDNA in rapidly replicating cells like the basal layer of the epidermis^28^ or in B cells^29^. Here, we have expressed K320E-Twinkle in dopamine neurons using DAT-cre expression as specific driver. Surprisingly, these animals do not develop a PD-like motor phenotype although 70% of SN dopamine neurons have perished at an advanced age of 20 months. This is due to compensatory branching of axon terminals in the dorsal striatum by the surviving SN dopamine neuron population, which enables normal dopamine signalling. Thus, we show that a small population of these neurons is able to adapt to physiologically occuring, harmful mtDNA mutations and to compensate severe neurodegeneration.

## Results

### Severe degeneration of dopaminergic midbrain neurons in K320E-Twinkle^DaN^ mice

To investigate the impact of slowly accumulating mtDNA alterations on dopaminergic neurons *in vivo*, mice harbouring a mutant variant of the mitochondrial helicase Twinkle (K320E, ^26^) were crossed with mice expressing Cre recombinase under control of the dopamine transporter promoter (DAT; Fig. 1a). Expression of K320E-Twinkle was confirmed by immunofluorescent detection of GFP, cloned downstream (Fig. 1b, Extended Data Fig. 1a). ∼98% of TH-positive neurons in the SN and VTA expressed GFP in 5-month-old mice (Fig. 1c). Conversely, ∼98% of GFP-positive neurons expressed TH, showing the specificity and effectiveness of Cre-mediated recombination in K320E-Twinkle^DaN^ dopamine neurons. While no difference in the total amount of TH-positive dopamine neurons was observed at 5 months of age, 10-month-old K320E-Twinkle^DaN^ mice revealed a significant loss of dopaminergic neurons (Fig. 1d, e). Mice carrying two copies of K320E-Twinkle (K320E/K320E^DaN^) presented a higher degree of neuron death (∼57%) than animals with only one copy (∼33%, K320E/Wt^DaN^). At the well-advanced age of 20 months, neurodegeneration was even more pronounced: K320E/K320E^DaN^ mice had lost ∼77% and K320E/Wt^DaN^ mice ∼41% of dopamine neurons. To examine whether the loss of neurons in K320E-Twinkle^DaN^ mice was restricted to a specific midbrain region, dopamine neurons were assigned to the SN and VTA, respectively. Neurons were lost to a rather similar extent in heterozygous (∼33%) and homozygous K320E-Twinkle^DaN^ mice (∼57%) at 10 months of age (Fig. 1f). From 10 to 20 months, the amount of dopamine neurons did not further change in K320E/Wt^DaN^ mice (for time line and statistics, see Extended Data Fig. 1b). In K320E/K320E^DaN^ mice, dopamine neurons further decreased in the VTA (from ∼55% to ∼84% loss), while there was no additional significant neuron loss in the SN (∼59% to ∼70% loss, Fig. 1f and Extended Data Fig. 1b).

**Figure 1.**
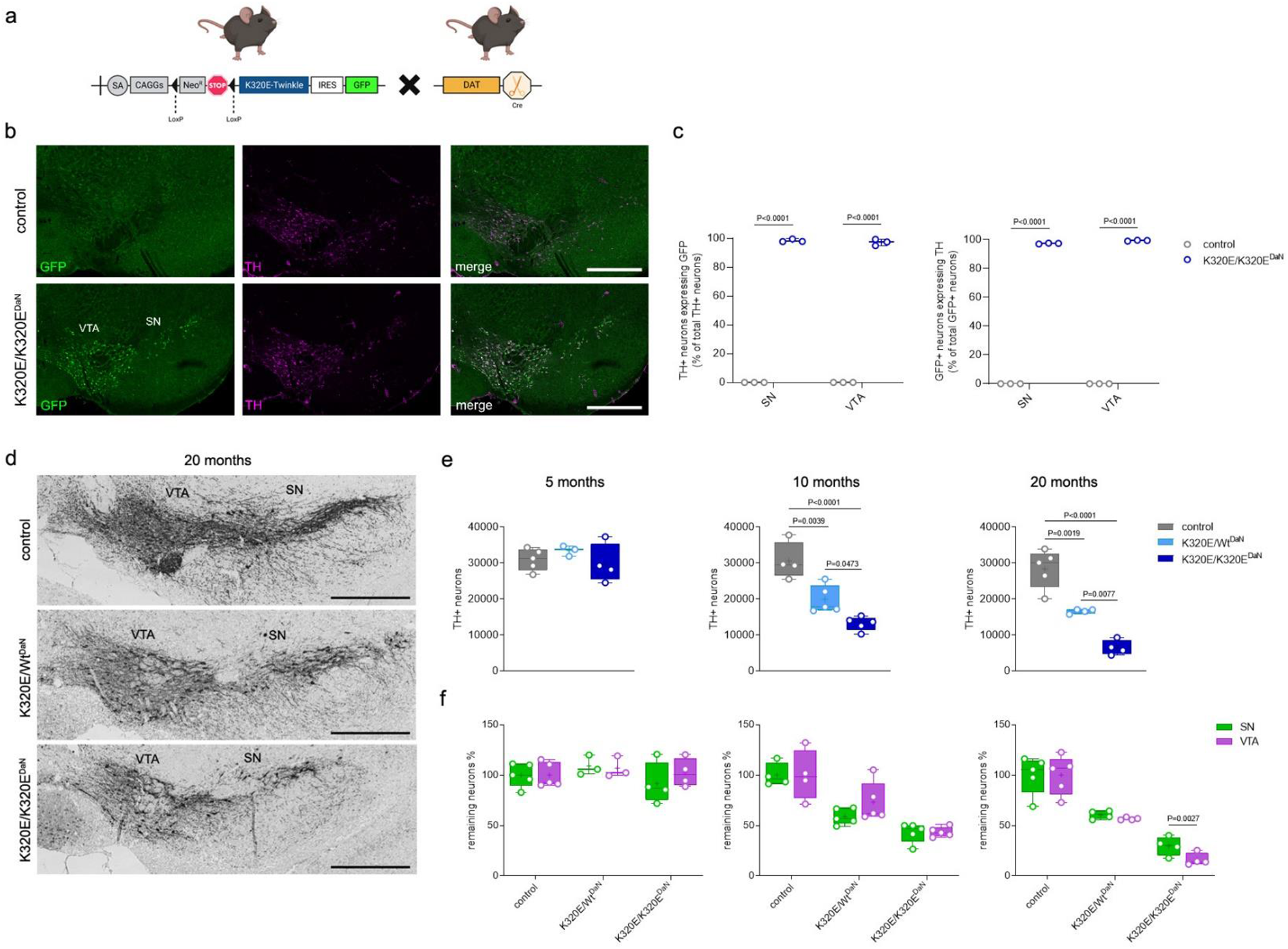
Age-dependent decline in SN and VTA dopamine neurons following K320E-Twinkle expression. **a**, Schematic of the generation of K320E-Twinkle^DaN^ mice. **b**, Representative images of tyrosine hydroxylase (TH) and GFP immunostaining in the midbrain of 5-month-old control and homozygous K320E-Twinkle^DaN^ mice; scale bar, 500 µm. **c**, Quantification of TH-positive cells expressing GFP as well as GFP-positive cells expressing TH (3 distinct Bregma levels from 3 animals per group). **d**, TH immunohistochemistry of coronal midbrain sections of 20-month-old control, K320E/Wt^DaN^, and K320E/K320E^DaN^ mice; scale bar, 500 µm. **e**, Stereological quantification of midbrain dopamine neurons and **f**, relative remaining neurons in the SN and VTA (5 months, n = 3 5; 10 months, n = 4 5; 20 months, n = 4 5). P values were calculated using one-way (e) or two-way ANOVA (c, f) with

### Neurodegeneration is preceded by reduced levels and activity of mitochondrial complex IV

To evaluate mitochondrial respiratory chain abundance in dopamine neurons upon K320E-Twinkle expression, we performed quantitative analysis of the mtDNA encoded subunit I assembled into cytochrome c oxidase (COX; complex IV) using immunofluorescent staining (Fig. 2a, representative micrographs display SN neurons). At 5 months of age, before the onset of neurodegeneration, COXI levels were significantly decreased in both SN and VTA dopamine neurons of hetero-and homozygous K320E-Twinkle^DaN^ mice (Fig. 2b). Noteworthy, intensity of COXI was not uniformly affected in all investigated neurons: While the mean COXI level was reduced by ∼36% (K320E/Wt^DaN^) and ∼49% (K320E/K320E^DaN^), single cell analysis revealed different populations of mutant dopamine neurons, including cells with much lower (up to ∼70%) as well as normal COXI levels when compared to controls (Fig. 2b). Also, reduced levels of subunit NDUFB11, assembled into complex I, were detected in both SN and VTA dopaminergic neurons of 5 months old K320E-Twinkle^DaN^ mice (Extended Data Fig. 2a). In addition, we investigated COX enzymatic activity by histochemical staining of frozen sections. Blue, COX-deficient cells were detectable in both SN and VTA of K320E-Twinkle^DaN^ mice at 5 months (Fig. 2c) and 10 months of age (Extended Data Fig. 2b-c). At 20 months, remaining neurons revealed normal COXI levels (Extended Data Fig. 2e). Accordingly, only few cells with deficient COX activity were detectable (Extended Data Fig. 2d).

**Figure 2.**
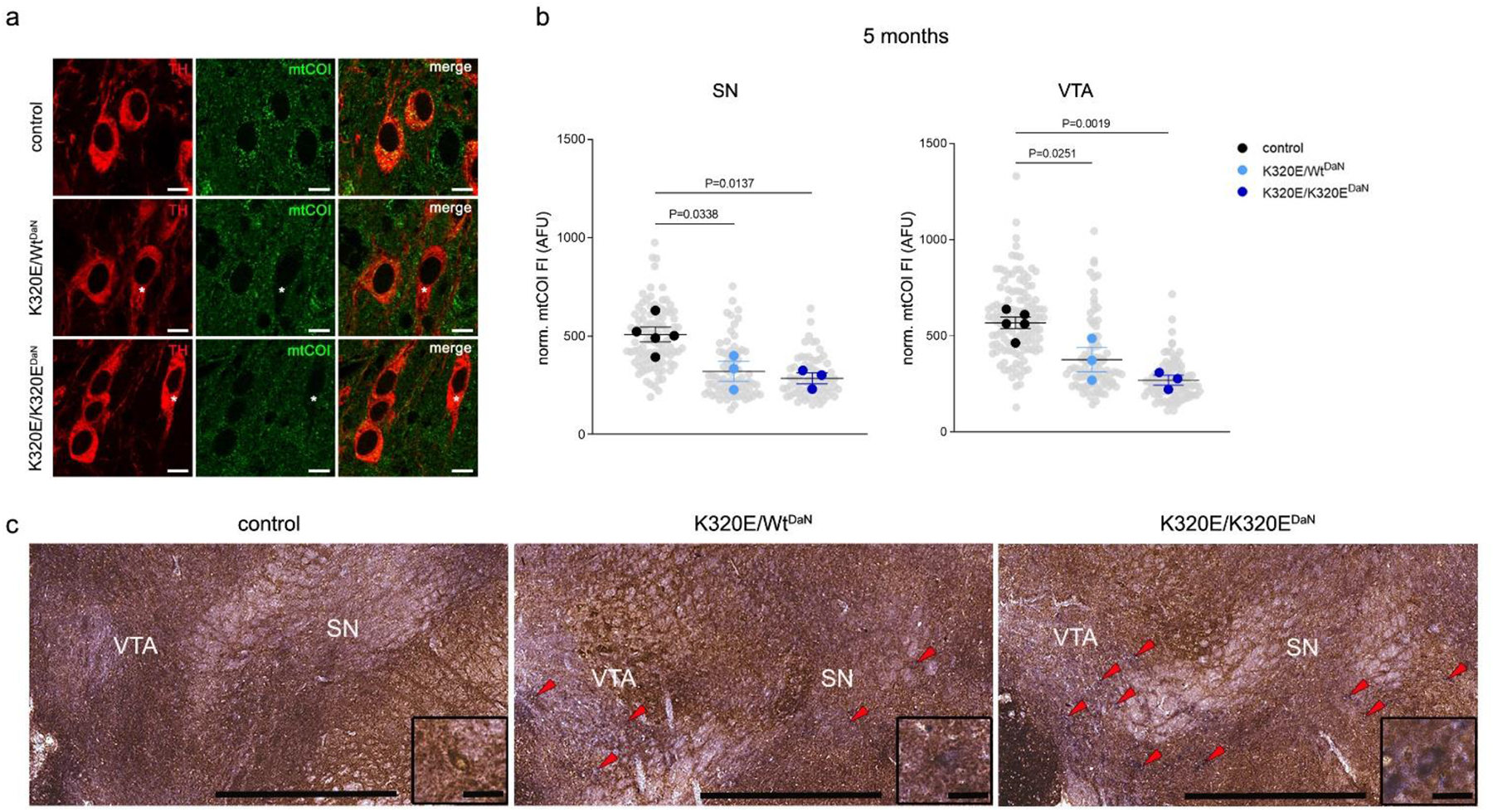
Reduced expression and activity of cytochrome c oxidase before the onset of neurodegeneration. **a**, Representative x63 confocal images of TH and mtCOI immunostaining in the SN of 5-month-old control, K320E/Wt^DaN^, and K320E/K320E^DaN^ mice; scale bar, 10 µm. **b**, Normalized mean fluorescent intensity (FI) of mtCOI in SN and VTA dopamine neurons (69-96 SN neurons, 88 125 VTA neurons, from 3 5 mice per group). P values were calculated using one-way ANOVA with Bonferroni’s correction for multiple comparison. Data are presented as superplots including mean ± s.e.m. **c**, Representative images of a dual staining for cytochrome c oxidase (COX) and succinate dehydrogenase (SDH) enzyme activity revealing blue, COX-deficient cells in the SN and VTA (red arrowheads) of 5-month-old K320E/Wt^DaN^ and K320E/K320E^DaN^ mice; scale bar, 500 µm and 20 µm, respectively.

### SN dopamine neurons accumulate mtDNA single nucleotide variants following K320E-Twinkle expression

With increasing age, SN dopamine neurons accumulated mtDNA indels in both control and K320E/K320^DaN^ mice (Fig. 3a). Expression of K320E-Twinkle did not change the numbers of deletions and duplications relative to control mice (Fig. 3b). We next examined the burden of mitochondrial single nucleotide variants (mtSNVs) of potential somatic origin (heteroplasmy fraction, HF, 1-5%) or after clonal expansion (HF 5-90%). Pooled SN dopamine neurons from 5-month-old K320E/K320E^DaN^ mice revealed a higher cumulative frequency of somatic mtSNVs compared to control mice and K320E/K320E^DaN^ animals at 20 months (Fig. 3c). In addition, the cumulative frequency of clonally expanded mtSNVs in 5-month-old K320E/K320E^DaN^ mice was comparable to the mtSNVs frequency of their 20-month-old counterparts, which suggests an early acquisition of mtSNVs resulting in a state of accelerated mtDNA ageing. mtSNV accumulation was obviously compensated by a pronounced increase of mtDNA copy number at 5 months, while the 30% surviving mutant neurons showed equally high copy numbers compared to the intact pool of control neurons at 20 months (Fig. 3d).

**Figure 3.**
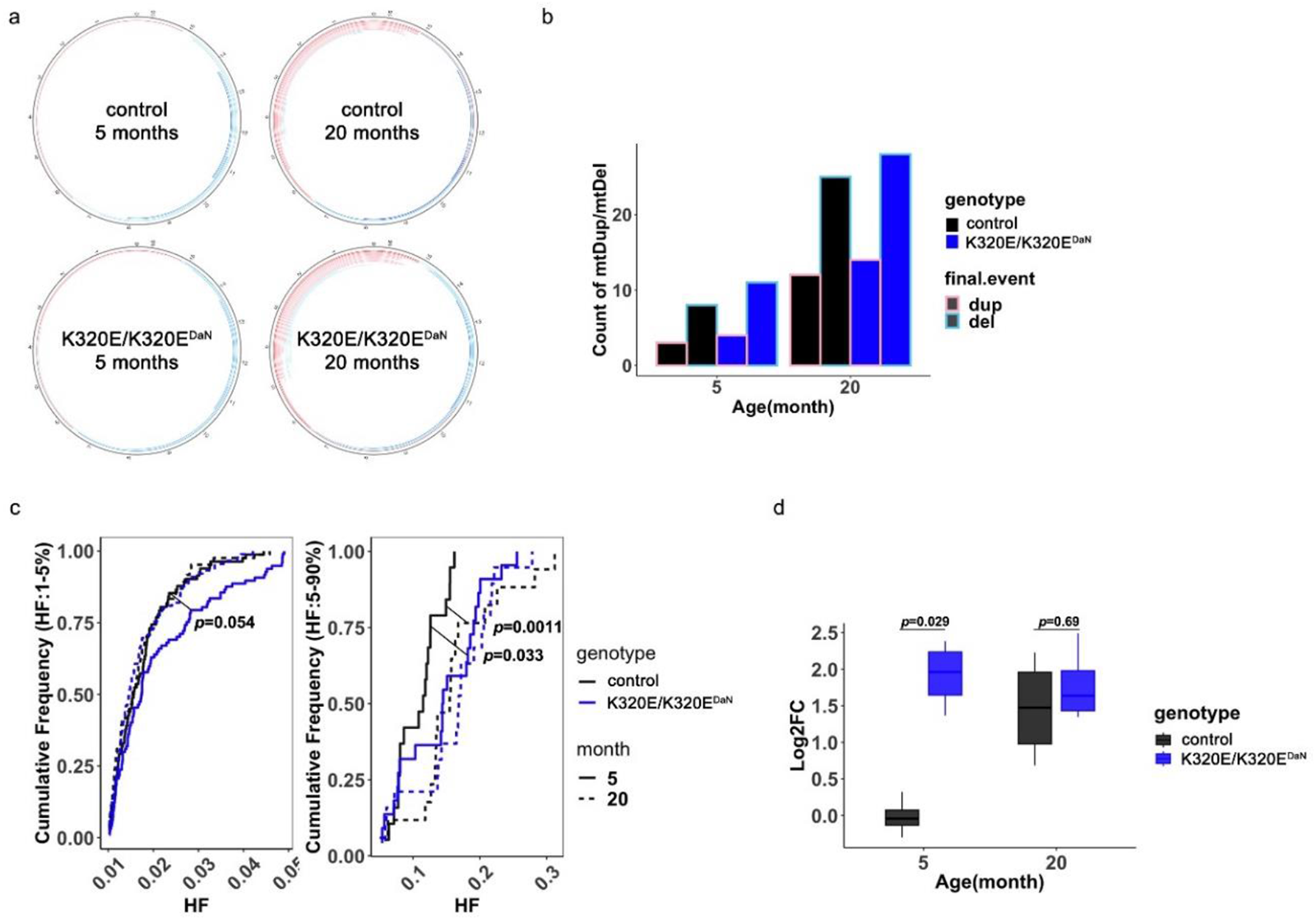
K320E-Twinkle expression causes mtDNA point mutations and copy number upregulation in SN dopamine neurons. **a**, **b**, **c**, **d**, mtDNA analysis of SN dopamine neurons isolated from control and homozygous K320E-Twinkle^DaN^ mice at 5 and 20 months. (**a**) Circular plots illustrating the distribution of mtDNA duplications (mtDup, pink) and deletions (mtDel, sky-blue) (3 mice per group). (**b**) Total counts of mtDup (pink outline) and mtDel (sky-blue outline) (3 mice per group). (**c**) Log2 fold change of mtDNA copy number, compared to control values at 5 months (4 mice per group). (**d**) Cumulative heteroplasmy fraction (HF) distribution of mitochondrial single nucleotide variants (mtSNVs) in control (black) and K320E/K320E^DaN^ (blue) animals at 5 months (solid line) and 20 months (dash line), between 1-5% HF (newly acquired mutations) and 5-90% HF (predicted clonal expanded mutations).

### Normal motor behaviour despite severe loss of SN dopamine neurons

Next, we explored the impact of SN neurodegeneration on motor function of K320E-Twinkle^DaN^ mice. Spontaneous motor activity was analysed using the open field BeamBreak test (Fig. 4a). Surprisingly, neither horizontal locomotion nor vertical rearing of K320E-Twinkle^DaN^ mice differed from control littermates at 20 months of age (Fig. 4b, c). In addition, motor coordination was examined by the RotaRod test (Extended Data Fig. 4a). 10- and 20-month-old K320E/K320E^DaN^ mice showed similar latency periods to fall from the rotating rod when compared to age-matched control mice (Extended Data Fig. 4b).

**Figure 4.**
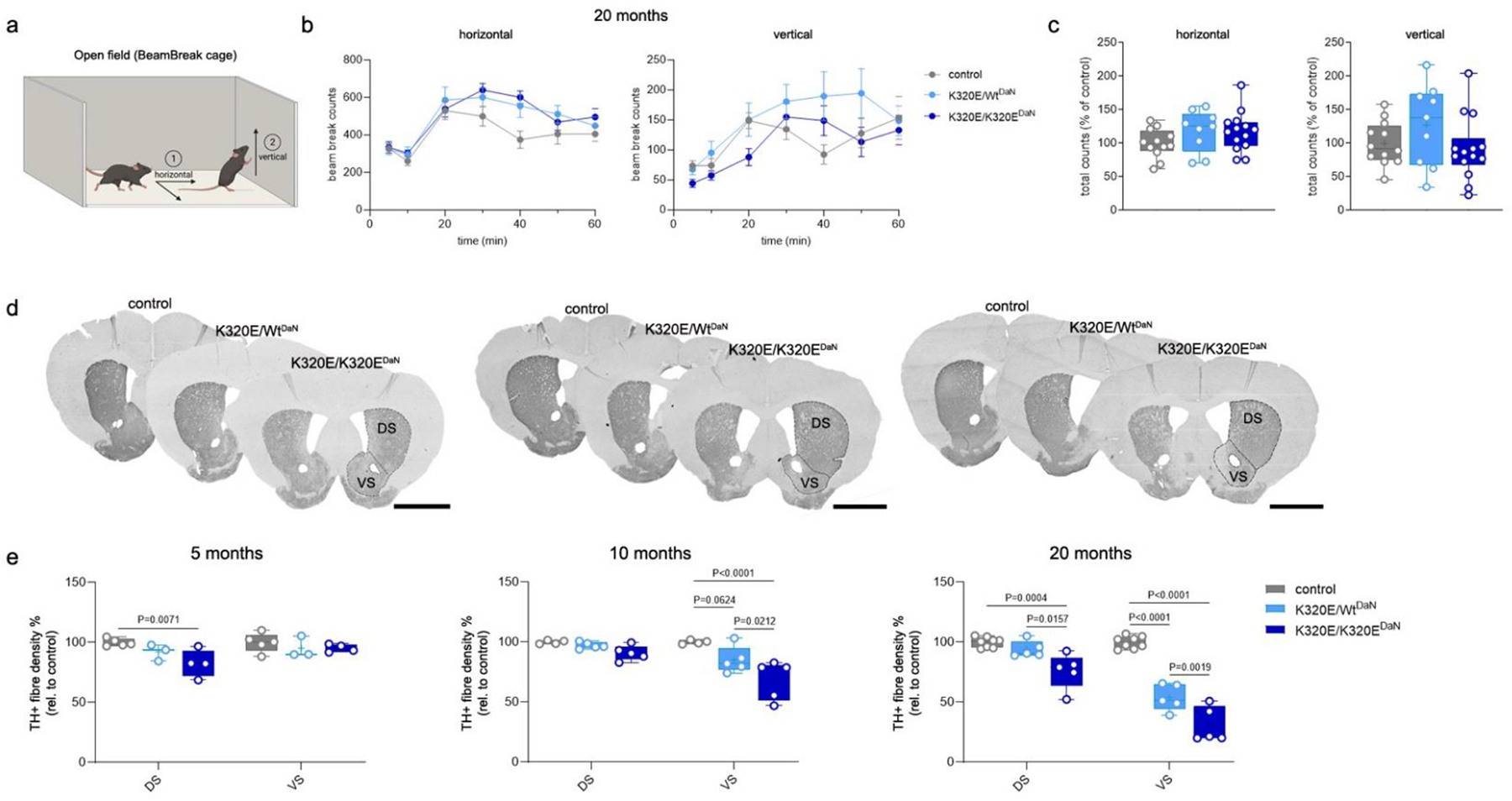
Normal motor behavior in K320E-Twinkle^DaN^ mice is associated with preserved dopaminergic terminals in the dorsal striatum. **a**, Schematic of the BeamBreak cage setup. **b**, Beam break counts for horizontal (locomotion) and vertical (rearing) movement of 20-month-old control, K320E/Wt^DaN^, and K320E/K320E^DaN^ mice over a period of 60 min (control, n = 12; K320E/Wt^DaN^, n = 9; K320E/K320E^DaN^, n = 14). **c**, Relative total counts after 60 min. **d**, Representative images of TH immunohistochemistry in the dorsal (DS) and ventral striatum (VS) at 5, 10, and 20 months; scale bar, 2 mm. **e**, Box plots presenting the quantitative analysis of TH-positive fibre density (5 months, n = 3 5; 10 months, n = 4 5; 20 months, n = 5 8). P values were calculated using two-way ANOVA with Bonferroni’s correction for multiple comparison. Data are presented as mean ± s.e.m.

### Dopaminergic innervation is preserved in the dorsal but not ventral striatum

Absent motor impairment in K320E-Twinkle^DaN^ mice suggests that, despite severe degeneration of SN dopamine neurons, sufficient levels of dopamine are released in the dorsal striatum to ensure normal motor function. To test this hypothesis, we first analysed the density of dopaminergic axon terminals in the striatum by TH immunohistochemistry (Fig. 4d). TH levels were determined in the dorsal and ventral striatum, which harbour axonal projections of dopaminergic neurons from SN and VTA, respectively. At 5 months, TH signal did not differ between K320E/Wt^DaN^ mice and control littermates (Fig. 4e), while K320E/K320E^DaN^ mice revealed a slight reduction of TH-positive fibres in the dorsal striatum (∼17%), whereas the ventral striatum was unchanged. Interestingly, similarly low TH levels were detected in the dorsal striatum for both heterozygous and homozygous K320E-Twinkle^DaN^ mice at 10 months of age. At that age, dopaminergic SN neurons were already lost by ∼41% (K320E/Wt^DaN^) and ∼59% (K320E/K320E^DaN^), respectively (Fig. 1). In contrast, TH-positive fibre density in the ventral striatum decreased by ∼15% in heterozygous and by ∼30% in homozygous K320E-Twinkle^DaN^ animals. The discrepancy in TH immunoreactivity between the dorsal and ventral striatum continued to grow after 20 months of age: Whereas TH-positive fibre density in the ventral striatum was decreased by ∼46% (K320E/Wt^DaN^) and ∼67% (K320E/K320E^DaN^), respectively, TH immunoreactivity in the dorsal striatum was preserved at ∼94% in heterozygous and ∼77% in homozygous K320E-Twinkle mice. Similar results were obtained by DAT immunohistochemistry (Extended Data Fig. 4c-g). Next, we tested if the loss of the mesolimbic dopamine system in K320E-Twinkle^DaN^ mice was associated with abnormal reward and social behaviour. While K320E-Twinkle^DaN^ animals showed normal immobility times during the tail suspension test (Extended Data Fig. 4h-i), they consumed significantly lower amounts of sucrose water in the sucrose preference test (Extended Data Fig. 4j, k). Moreover, K320E/K320E^DaN^ mice showed a tendency for increased social affiliation but normal behavior when exposed to new social stimuli in the three chamber test (Extended Data Fig. 4l-o).

### Decreased dopamine synthesis capacity but normal transients of striatal dopamine release

Next, we explored the *in vivo* dopamine synthesis and storage capacity in the striatum by positron emission tomography (PET) using the radioligand [^18^F]FDOPA (Fig. 5a). While 20-month-old control mice showed pronounced [^18^F]FDOPA uptake, it was considerably reduced in K302E/K320E^DaN^ mice (Fig. 5b). VOI analysis revealed a mean SUVR_CB_ of 1.12 ± 0.02 in controls and 1.06 ± 0.01 in K320E/K320E^DaN^ mice. Noteworthy, dopaminergic innervation of the cerebellum is minimal and dopamine binding sites are ∼10 times less abundant than in the striatum^30^. Cerebellar [^18^F]FDOPA uptake thus mostly reflects unspecific accumulation and was used as background signal for intensity normalization. Since cerebellar [^18^F]FDOPA uptake was set to 1, the striatal [^18^F]FDOPA uptake in K320E/K320E^DaN^ mice can be interpreted as a ∼50% reduction compared to control animals. PET tracers reflecting presynaptic density ([^18^F]MNI1126) and general neuronal metabolic activity ([^18^F]FDG) revealed no difference between control and K320E/K320E^DaN^ mice (Extended Data Fig. 5a, b).

**Figure 5.**
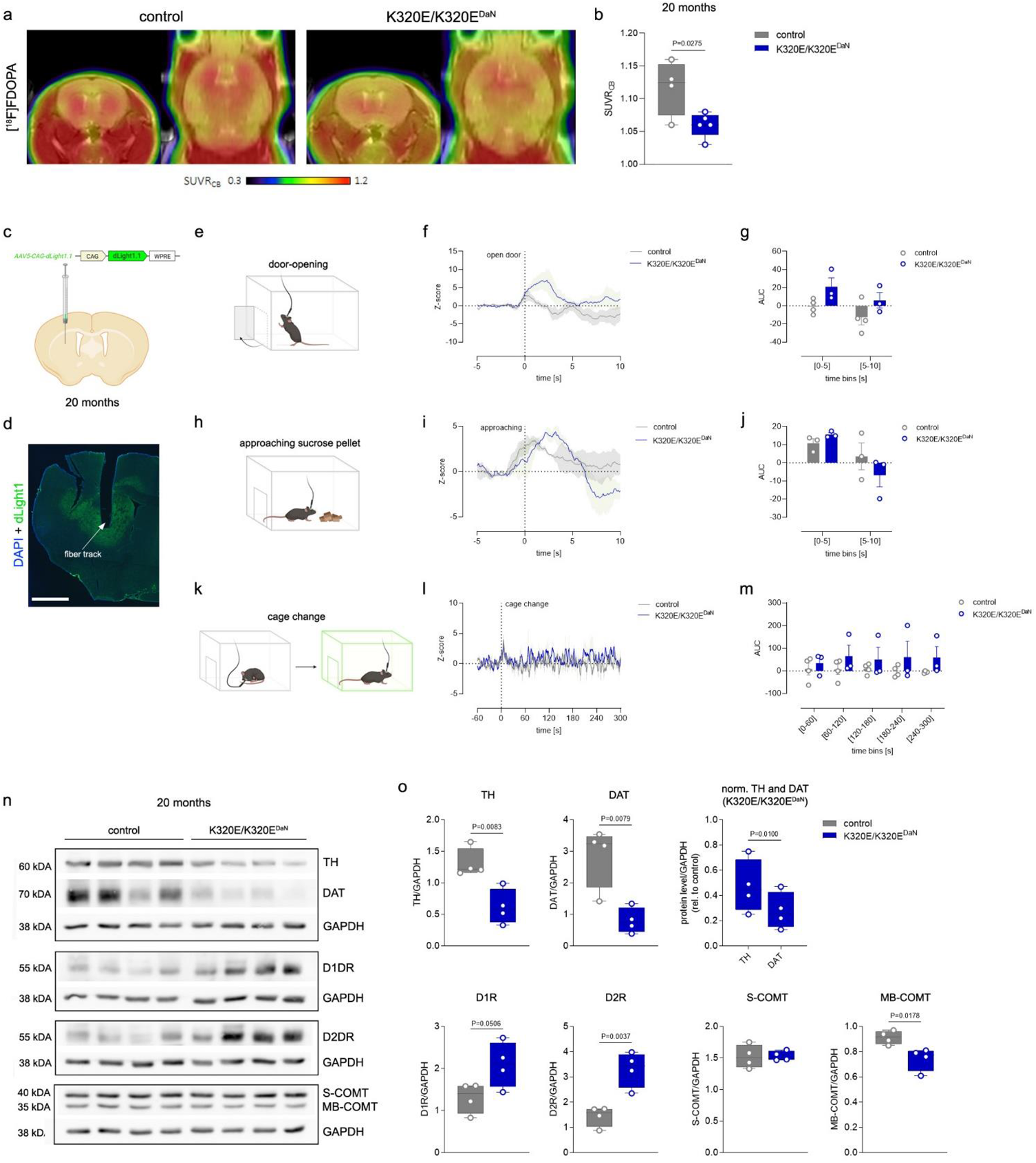
Compensatory dopamine metabolism in the striatum of K320E/K320E^DaN^ mice. **a**, Averaged PET scan images presenting the uptake of [18F]FDOPA by 20-month-old control and K320E/K320E^DaN^ mice. **b**, PET scan quantification showing the standardized uptake value ratios (SUVRCB) for [18F]FDOPA in the striatum. P value was calculated using unpaired two-tailed Student’s t test. **c**, Schematic of AAV5 CAG dLight1.1 injection into the dorsal striatum. **d**, Expression of dLight1.1 in the dorsal striatum with track of the applied optical fiber; scale bar, 1 mm. **e**, Schematic showing sudden door-opening of the experimental cage. **f**, Dopamine response to opening of the experimental setup’s door (t=0) represented as Z-score relative to baseline (t = [-5-1]). **g**, Area under the curve (AUC) of the dopamine response to sudden opening of the experimental cage’s door, calculated for 5 sec bins (control, n = 4; K320E/K320EDaN, n = 3). **h**, Schematic showing the offer of sucrose pellets. **i**, Dopamine response to sucrose pellet (t=0) represented as Z-score relative to baseline (t = [-5-1]). **j**, AUC of the dopamine response to sucrose pellet, calculated for 5 sec bins (control, n = 4; K320E/K320E^DaN^, n = 3). **k**, Schematic showing the change of the experimental cage. **l**, Dopamine dynamics while mice were exposed to a new cage (t=0), represented as Z-score relative to baseline (t = [-60-1]). **m**, AUC of the dopamine response to the fresh cage, calculated for 60 sec bins (control, n = 4; K320E/K320E^DaN^, n = 3). **n**, **o** Western blot analysis and quantification of proteins involved in dopamine metabolism in the striatum of control and K320E/K320E^DaN^ mice at 20 months. P value indicating significant difference between TH and DAT levels was calculated using paired two-tailed Student’s t test. Residual P values were calculated using unpaired two-tailed Student’s t test. Data are presented as mean ± s.e.m.

It has been shown that in striatal dopamine axons only ∼30% of varicosities contain active zone-like sites, which can be identified by the expression of the scaffolding protein bassoon^31^. Co-staining for TH and bassoon showed similar numbers of bassoon expressing varicosities in TH-labelled dopamine axon terminals in both genotypes (Extended Data Fig. 5c-f), excluding an increased recruitment of active zone-like sites as a mechanism for sufficient dopamine release.

To examine striatal dopamine release *in vivo*, we next performed fiber photometry using the genetically encoded extracellular dopamine optical sensor dLight1.1^32^. An AAV encoding the dLight1.1 construct was unilaterally injected into the dorsal striatum of 20-month-old mice (Fig. 5c), followed by implantation of an optical fiber for induction and collection of the dLight1.1 signal (Fig. 5d). Dopamine release was triggered in awake, freely behaving mice by three salient stimuli, which are well characterized to evoke dopaminergic activity: sudden door-opening of the experimental cage (Fig. 5e), delivery of a palatable sucrose pellet (Fig. 5h) and change to a new cage (Fig. 5k)^33^. K320E/K320E^DaN^ mice revealed attenuated raw dLight1.1 signals compared to control animals, suggesting lower basal and absolute striatal dopamine levels (Extended Data Fig. 5g). Importantly, however, the relative dopamine response was indistinguishable in both dynamics and intensity between K320E/K320E^DaN^ and control mice following all three stimuli (Fig. 5e-m). Altogether, dopamine photometry analyses revealed that K320E/K320E^DaN^ mice may have lower basal and absolute striatal dopamine levels, which is in agreement with our [^18^F]FDOPA results. However, the relative increase in striatal dopamine in response to different salient stimuli is unaffected.

Analysis of striatal protein levels revealed low TH and DAT, reflecting the cumulative loss of dopamine axon projections in the combined dorsal and ventral striatum (Fig. 5h, i). Noteworthy, relative levels of DAT were more reduced than TH in K320E/K320E^DaN^. K320E/K320E^DaN^ showed enhanced levels of both dopamine 1 (D1R) and 2 receptors (D2R), which was accompanied by decreased levels of catechol-o-methyltransferase (COMT), which is involved in dopamine degradation and predominantly represented in the brain by its membrane-bound isoform (MB-COMT)^34^. There were no changes in protein levels of MAO-A (Extended Data Fig. 5h, i), which was recently shown to be the monoamine oxidase mainly contributing to dopamine degradation^35^. Levels of MAO-B, conversely, were increased but this was not due to an enhanced presence of astrocytes, where MAO-B is primarily localized^36, 37^, according to the astrocytic marker glial fibrillary acidic protein (GFAP).

### Transcriptome of surviving SN dopamine neurons reveals upregulated pathways stimulating axon guidance, calcium signalling and neuroactive ligand-receptor interaction

To unmask changes in gene expression following K320E expression, cDNA from control and K320E/K320E^DaN^ SN dopamine neurons was sequenced at 5 and 20 months of age (Fig. 6a). SN neurons showing TH immunoreactivity were precisely isolated by laser capture microscopy^38^ (Extended Data Fig. 6a-c) and revealed gene expression enrichment characteristic for dopamine neurons (Fig. 6b). Genes differentially expressed in SN dopamine neurons of 5-month-(Extended Data Fig. 6e, g) and 20-month-old K320E/K320E^DaN^ mice (Fig. 6c and Extended Data Fig. 6d) were clustered by pathway and process enrichment analysis. Among the most powerfully represented pathways, surviving SN dopamine neurons revealed upregulation of calcium signalling and neuroactive ligand-receptor interaction, as well as axon guidance and regulation of actin cytoskeleton (Fig. 6c). Protein-protein interaction analysis of upregulated genes identified calcium and acetylcholine signalling, Ras signalling, and axon guidance as the most strongly pronounced networks (Fig. 6d). Axon guidance was also identified as upregulated pathway and pronounced network in SN dopamine neurons of 5-month-old K320E/K320E^DaN^ animals (Extended Data Fig. 6e. f). Due to the striking upregulation of receptors for factors modulating axon growth in surviving SN dopamine neurons, we examined gene expression of neurotrophic factors and guidance molecules in striatal homogenates from 20-month-old K320E/K320E^DaN^ mice. While there was no difference in mRNA levels for the conventional neurotrophic factors *Ngf*, *Bdnf* and *Gdnf*, K320E/K320E^DaN^ mice revealed upregulation of *Netrin-1* and *Ephrin-A2* (Fig. 6e), which have both been reported to positively influence the innervation of the dorsal striatum by SN dopamine neurons^39, 40^. This was accompanied by downregulation of *Sema3A* and *Slit-2*, which in turn, are linked to the inhibition of axon branching^41–43^.

**Figure 6.**
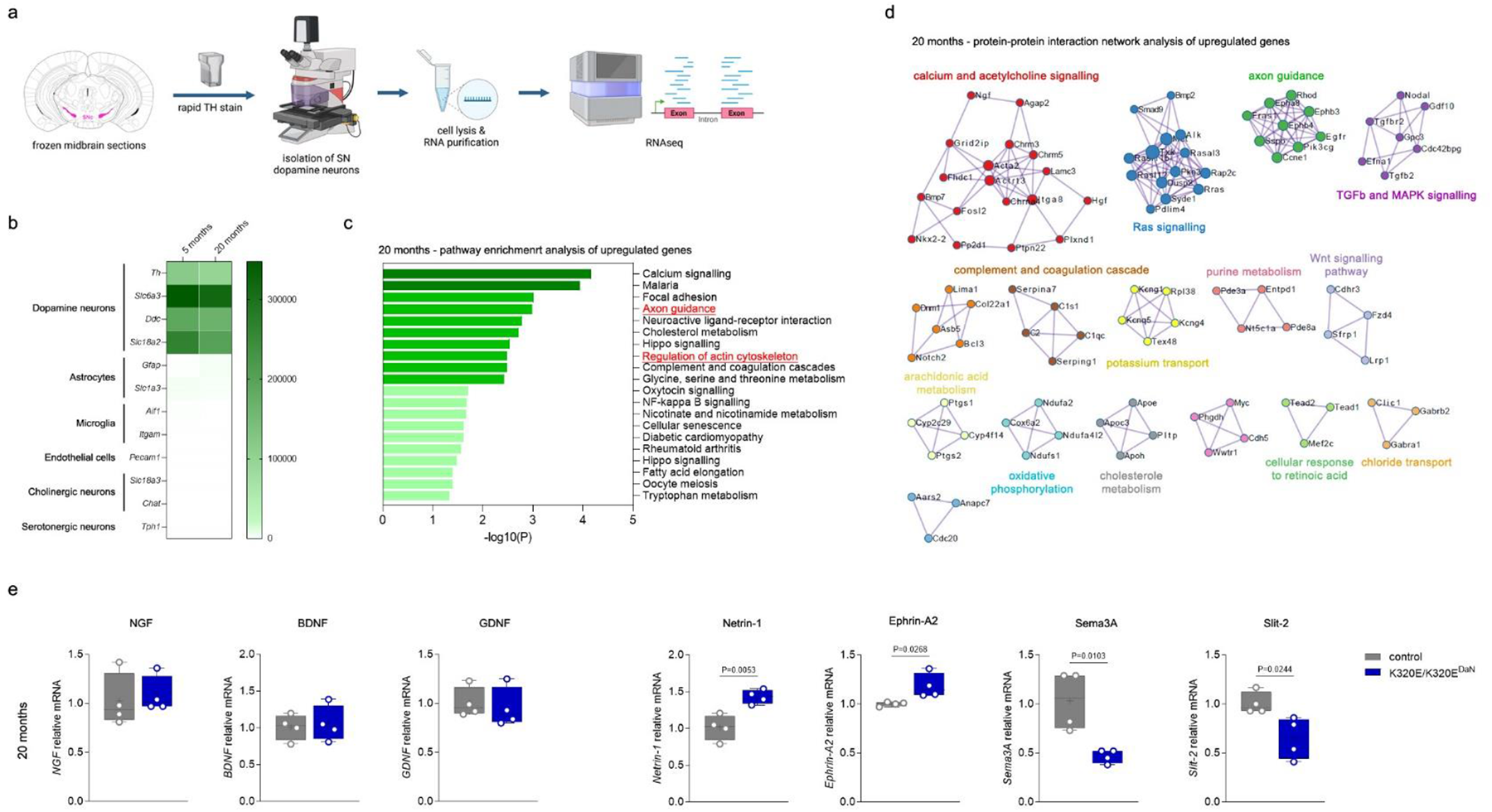
RNAseq of SN dopamine neurons uncovers upregulation pathways involved in axon guidance, calcium signalling and neuroactive ligand-receptor interaction in K320E/K320EDaN mice. **a**, Workflow of RNAseq following laser capture microscopy of SN dopamine neurons. **b**, Heat map showing the amount of total read counts for genes expressed by distinct cell types in laser-captured SN neurons at 5 and 20 months. **c**, Bar graph illustrating enriched upregulated pathways in SN dopamine neurons of 20-month-old K320E/K320E^DaN^ mice. The top 20 most statistically significant clusters are shown; -log(10)P is the P value in negative log base 10. **d**, Network visualization of enriched protein-protein interaction for upregulated genes in 20-month-old K320E/K320E^DaN^ mice. **e**, qPCR mRNA quantification of axon guidance molecules in the striatum of 20-month-old K320E/K320E^DaN^ and control mice. Values were normalized to Gapdh mRNA (n = 4). P values were calculated using unpaired two-tailed Student’s t test. Data are presented as mean ± s.e.m.

### Surviving SN dopamine neurons preserve dorsal striatum innervation

Our results indicate that the remaining population of SN dopamine neurons in K320E/K320E^DaN^ mice preserve striatal dopamine supply by compensatory axonal sprouting despite the presence of K320E-Twinkle. Although we have shown that almost 100% of SN dopamine neurons expressed the construct before the onset of neuron loss (Fig. 1b, c), a small proportion of the surviving cells could still lack the mutant Twinkle. Noteworthy, previous studies reported on the existence of TH-positive neurons with little or no DAT expression^44, 45^, which in our case, would affect the efficiency of DAT-Cre-mediated recombination. Immunofluorescent detection of TH and DAT, however, revealed a high level of colocalization in remaining SN neurons of 20-month-old K320E/K320E^DaN^ mice (Extended Data Fig. 7a-c). To collectively confirm K320E expression and the compensatory axonal sprouting of surviving SN dopamine neurons in K320E/K320E^DaN^ mice, we performed anterograde viral tracing, using an AAV that expresses EGFP under control of the Cre/loxP system through a modified Flex switch (AAV-FLEX-EGFP; Fig. 7a). AAV-FLEX-EGFP was unilaterally injected into the SN of 20-month-old K320E/K320E^DaN^ mice and corresponding control animals expressing DAT-Cre (Fig. 7b). Viral injection led to a robust expression of EGFP in TH-labelled dopamine midbrain neurons (Fig. 7c). The infection efficiency of TH-labelled dopamine neurons in the SN was about ∼90% for both K320E/K320E^DaN^ and control animals and significantly higher than in the VTA (∼45%; Fig. 7d). Conversely, ∼99% of EGFP-positive neurons expressed TH, which excludes potential off-target viral infection (Extended Data Fig. 7d). The number of remaining midbrain neurons in K320E/K320E^DaN^ animals that express TH and EGFP was consequently reduced to a similar extent (Fig. 7e and Extended Data Fig. 7e). In line with the high infection efficiency of SN dopamine neurons, analysis of the axon projection area revealed EGFP expression preferentially in the dorsal striatum for both K320E/K320E^DaN^ and control mice (Fig. 7f, g). Non-injected hemispheres were devoid of EGFP signal (Extended Data Fig. 7f). In comparison to the dorsal striatum of control animals, EGFP-positive fibre density of K320E/K320E^DaN^ mice was decreased by ∼59% (Fig. 7g), but importantly, matched with TH-labelled axon terminals by ∼91% (Fig. 7h).

**Figure 7.**
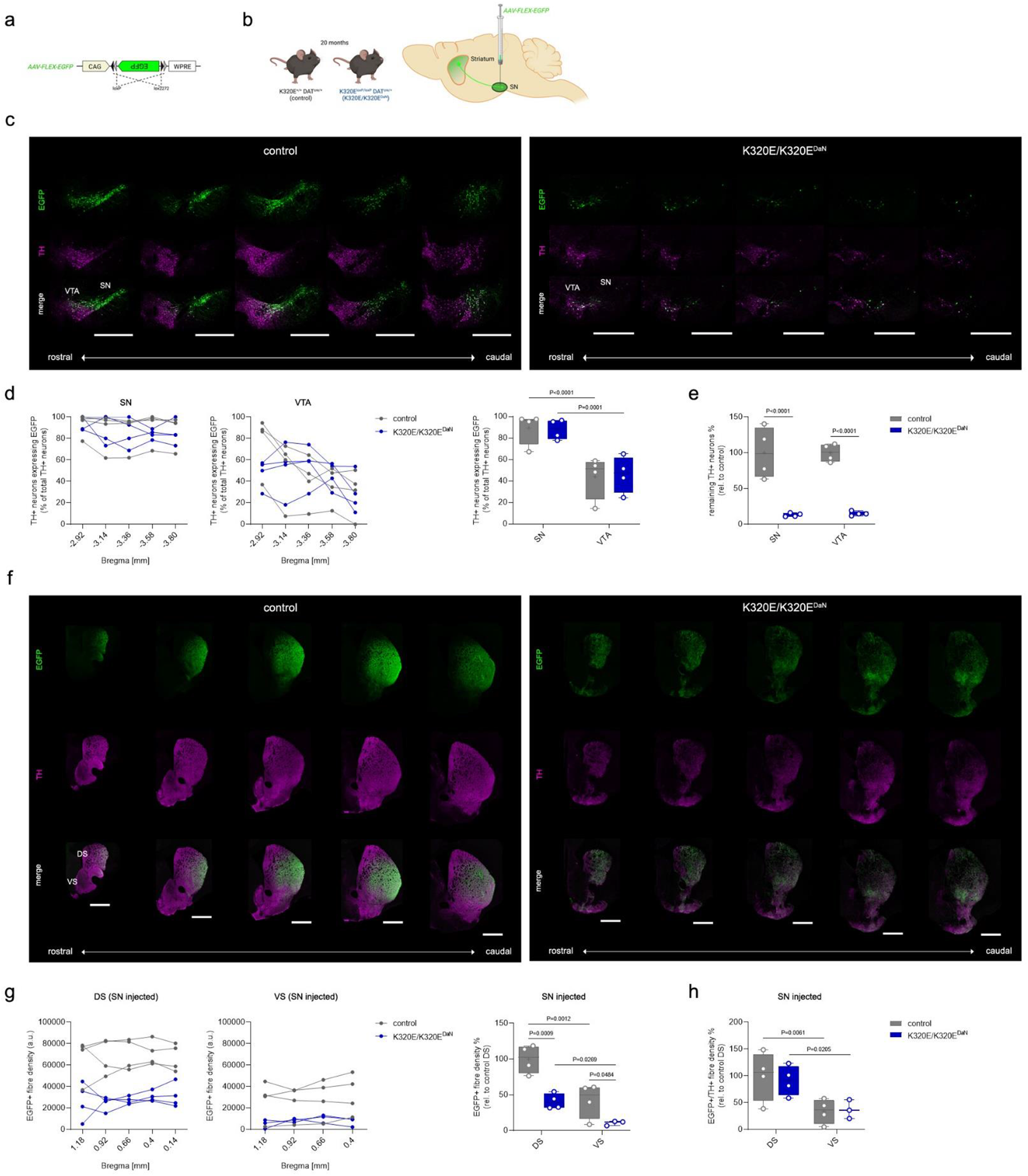
Preserved dopaminergic fibres in the dorsal striatum originate from the SN. **a**, AAV strain used for CAG-driven EGFP expression in the presence of Cre. **b**, Schematic of anterograde viral tracing of SN dopamine neurons in 20-month-old control and K320E/K320E^DaN^ mice. **c**, Representative immunofluorescent images showing EGFP expression in SN and VTA dopamine neurons along the rostro-caudal axis; scale bar, 1 mm. **d**, Quantification of TH positive neurons expressing EGFP. **e**, Relative remaining TH-positive neurons in the SN and VTA (5 distinct Bregma levels from 4 animals per group). **f**, Representative immunofluorescent images showing EGFP expression in the dorsal (DS) and ventral striatum (VS) along the rostro-caudal axis upon viral injection into the SN; scale bar, 1 mm. **g**, Quantification of EGFP-positive fibre density in the striatum. **h**, Ratio between EGFP-and TH-positive fibre density (DS: 5 distinct Bregma levels from 4 animals per group, VS: 4 distinct Bregma levels from 4 animals per group). P values were calculated using two-way ANOVA with Bonferroni’s correction for multiple comparison. Data are presented as mean ± s.e.m.

## Discussion

In idiopathic PD, a high mtDNA mutation load ^22^ meets low wildtype copy numbers in dopamine neurons, in contrast to unaffected individuals in which copy number increases during ageing^25^, which renders impaired mtDNA homeostasis an important puzzle piece for age-dependent SN dopamine neuron loss. The K320E-Twinkle mutation has been shown to slow down mtDNA replication in dividing cells^46^ and to induce indels in non-dividing cells^26, 27^. Our data show that in SN dopamine neurons, neither deletions nor duplications of mtDNA were further increased by K320E expression. Instead K320E-Twinkle led to an early accumulation of SNVs before the onset of neurodegeneration, resulting in a state of accelerated mtDNA ageing, which has not been reported before. As a response, SN dopamine neurons from K320E/K320E^DaN^ mice revealed an early compensatory upregulation of mtDNA copy number to levels present in control animals after 20 months, similar to what is observed in healthy ageing humans, but not in idiopathic PD patients^25^. High levels of wild-type mtDNA have consistently been shown to counteract the consequences of mtDNA mutations^48–50^. However, the early accumulation of SNVs was associated with the reduction in respiratory chain complexes and caused degeneration of a large neuron population in both SN and VTA starting between 5 and 10 months. In contrast to pooled SN dopamine neurons from K320E/K320E^DaN^ mice at 5 months of age, the surviving population after 20 months showed normal ageing-associated levels of SNVs which was accompanied by normal respiratory chain function, suggesting efficient mitochondrial quality control. The E3 ubiquitin ligase Parkin regulates the clearance of dysfunctional mitochondria via mitophagy^51^. Mutations in the encoding *PARK2* gene cause autosomal recessive PD with early disease onset ^52^ and the ligase activity of Parkin is assumed to be affected in idiopathic cases as well^53^. We therefore tested the hypothesis if in absence of Parkin, the phenotype of K320E/K320E^DaN^ animals would get worse and motor deficits might emerge. The lack of Parkin did neither change the number of remaining dopamine neurons nor the density of dopamine axon terminals nor motor behaviour (Extended Data Fig. 8a-f). We hence conclude that in surviving SN dopamine neurons of K302E/K320E^DaN^ mice, mitochondrial quality control occurs in a Parkin-independent manner, which is in line with previous studies questioning the relevance of Parkin for the survival of dopamine neurons in rodents^54, 55^. Besides the variety of mitophagy pathways that can act in absence of Parkin^56^, our group has recently described a pathway for the selective turnover of mutation-bearing mtDNA involving VPS35^57^, mutations in which are linked to autosomal dominant PD^58^.

Although ∼70% of SN dopamine neurons had perished, K320E/K320E^DaN^ mice did not show any signs of motor impairment at the late age of 20 months, which is in stark contrast to other genetically or chemically induced PD mouse models^59–65^. Additional immunohistochemical approaches as well as viral tracing confirmed the severe loss of dopamine neurons (Fig. 7e and Extended Data Fig. 7a-c, e) and excluded downregulation of TH as the reason. The normal motor behaviour is explained by the maintenance of ∼75% of dopamine axon terminals in the dorsal striatum. In the ventral striatum, on the other hand, only ∼30% of dopamine projections originating from the VTA remained, which caused abnormalities in reward and social behaviour of K320E/K320E^DaN^ animals. The largely preserved dopaminergic innervation in the dorsal striatum of K320E/K320E^DaN^ mice was accompanied by a ∼50% reduction in [^18^F]FDOPA uptake in the entire striatum, indicating a decreased capacity of L-DOPA conversion to dopamine and/or its storage^66, 67^. However, normal uptake of [^18^F]MNI1126 and [^18^F]FDG, suggests a compensatory upregulation of presynaptic density and, consequently, neuronal metabolic activity. Nevertheless, both tracers are not specific for dopamine axons and the dopamine terminal fraction might be too small to see significant differences in their uptake of both tracers. In line with this, an analogous tracer to [^18^F]MNI1126 revealed no difference in the striatum of PD patients^68^. Since [^18^F]MNI1126 only detects synaptic vesicle glycoprotein 2A (SV2A)^69^, it is also possible that SV2A is not the predominant SV2 protein in dopamine axon terminals, as reported recently^70^. Finally, K320E/K320E^DaN^ mice did not show an upregulation of active-zone release sites at dopamine axon terminals.

Noteworthy, PET scanning was performed in anaesthetized animals. To analyse *in vivo* dopamine transients in freely behaving, awake K320E/K302E^DaN^ mice, we used the extracellular dopamine sensor dLight1.1 inserted into the dorsal striatum. In line with our [^18^F]FDOPA results, K320E/K320E^DaN^ mice showed lower baseline dopamine levels. Most importantly, however, the relative dopamine response following different tasks was indistinguishable from control animals. In addition, K320E/K320E^DaN^ mice show a compensatory upregulation of striatal D1Rs and D2Rs, indicating an increased sensitivity of medium spiny neurons, as well as downregulation of DAT and COMT, suggesting decreased breakdown. Upregulation of D2Rs^71^ and downregulation of DAT^6, 72, 73^ are phenomena also found in the early stage of idiopathic PD. In contrast to K320E/K320E^DaN^ mice, in PD patients this compensatory response is preceded by a ∼50-70% loss of dopamine axon projections in the striatum^4–7^.

Preserved dopamine axon terminals in the dorsal striatum of K320E/K320E^DaN^ mice are associated with the upregulation of pathways promoting axonal growth and branching in surviving SN dopamine neurons. We identified genes operating in Ras signalling, which is involved in nearly all stages of axiogenesis (reviewed by Hall et al.^74^), and TGFβ and MAPK signalling, promoting dopamine neuron survival and branching *in vitro* and *in vivo*^75, 76^. Among the upregulated genes involved in axon guidance we found *Pak4*, which has previously been shown to prevent degeneration of SN dopamine neurons and motor impairment in rat models of PD^77^. The upregulation of receptors for cues modulating axon growth, including ephrin, prompted us to also examine expression of genes for axon guidance molecules in the striatum. K320E/K320E^DaN^ mice showed elevated mRNA levels of *Netrin-1* and *Ephrin-A2*. Netrin-1 is known to promote axon growth and branching *in vitro*^41, 42^, and more importantly, it restored dopamine axon projections in the striatum of pharmacologically induced PD mouse models^39^. Ephrin-A signalling contributes to the guidance of dopamine projections towards the dorsal striatum during development^40^, a potential link for Ephrin-A2 to dopamine axon sprouting at advanced age, however, has not been reported before. K320E/K320E^DaN^ mice further showed downregulation of striatal *Sema3A* and *Slit-2*, which are both reported to inhibit axonal growth and branching^41, 42, 78, 79^. While gene expression analysis in PD patients suggests a rather chemorepulsive environment which might contribute to the denervation of striatal dopamine projections^80^, transcriptional changes in both the striatum and surviving SN dopamine neurons of K320E/K320E^DaN^ mice indicate predominant signalling towards the attraction and stabilization of dopamine axon terminals. The candidates presented here could hence be of relevance for the induction of dopamine axon sprouting, as supported by preclinical studies^39, 77^. Genome wide analyses of single nucleotide polymorphisms in PD patients showed axon guidance among the most significant pathways affected, including polymorphisms for the Netrin-1 receptor *DCC*, several Ephrin-A and -B receptors, as well as the Slit receptor *ROBO3*^81, 82^.

Intriguingly, calcium and acetylcholine signalling was upregulated as well in surviving SN dopamine neurons of K320E/K320E^DaN^ mice, which could not only have positive effects on axon growth^83–85^. The group of Pascal Kaeser recently presented a mechanism for striatal dopamine release supported by acetylcholine, enabling the local control of dopamine release independent of electrical input from the midbrain^86^. Regarding the low number of SN dopamine neurons left and the enhanced metabolic burden associated with axon sprouting^87^, it might thus be possible that normal dopamine release in K320E/K320E^DaN^ mice is additionally supported by local cholinergic transmission.

In line with the transcriptional adaptations, we finally showed by viral tracing that the preserved dopamine innervation in the dorsal striatum of 20-month-old K320E/K320E^DaN^ mice indeed came from the surviving population of SN neurons. Noteworthy, the proportion of remaining dopamine projections in the dorsal striatum was lower compared to our previous densitometric analyses using TH and DAT immunohistochemistry (Fig. 4d-e and Extended Data Fig. 4c-g). This was certainly due to the stereotaxic injection procedures (see Extended Data Fig. 7g), most likely into the dorsal striatum (data not shown), which are known to locally cause tissue damage^88, 89^. In the non-injected hemisphere, in turn, the density of dopamine fibres in the dorsal striatum of K320E/K320E^DaN^ mice was similar to the results of prior densitometric analyses. Importantly, TH-labelled axon terminals matched with the EGFP signal originating from SN dopamine neurons by ∼91%.

Together, our data show that following accelerating mtDNA mutagenesis in mice, a subset of SN dopamine neurons adapts to this insult and is able to compensate the severe reduction of the entire population at advanced age by axonal sprouting, enabling normal dopamine release and thus motor performance. Transcriptional changes in surviving SN dopamine neurons and its projection area draw further attention to adult axon branching induced by guidance molecules, including Netrin-1 and Ephrin-A2, that could be of particular importance for dopamine replacement therapies.

## Methods

### Animals

K320E-Twinkle^DaN^ and control mice were bred at the University of Cologne. Heterozygous (K320E/Wt^DaN^) and homozygous (K320E/K320E^DaN^) K320E-Twinkle^DaN^ animals were generated by crossing DAT-*cre* mice (Cre gene inserted upstream of the translation start codon in exon 2 of the DAT gene^65^ with K320E-Twinkle transgenic mice (point mutation K320E; Rosa26-Stop-construct; downstream GFP). DAT-*cre* mice had been kindly provided by Nils-Göran Larsson (Karolinska Institutet, Sweden). K320E-Twinkle animals had been generated previously by our group^26^. For the additional knockout (KO) of Parkin, K320E-Twinkle^DaN^ mice were crossed with Parkin KO animals, kindly provided by Olga Corti (Paris Brain Institute, Sorbonne University, France). Control mice were littermates harboring the K320E-Twinkle construct (heterozygous or homozygous) in absence of the Cre allele (except viral tracing experiment). Mice were kept in individually ventilated cages at 23 °C, 12:12 h light–dark cycle, with specified pathogen-free hygiene levels, free access to water and a regular chow diet ad libitum. All animals were regularly monitored for potential signs of pain and suffering. Breeding and experiments were conducted in agreement with European and German guidelines and approved by local authorities (LANUV, Landesamt für Natur, Umwelt und Verbraucherschutz Nordrhein-Westfalen, Germany; approval licenses: 84-02.04.2013-A141, 84-02.04.2016.A410, 81-02.04.2018-A210, 81-02.04.2021-A256). For all experiments, female and male animals were used. For behavioural assessments, group-housed animals were used to exclude the influence of social deprivation. Mice were genotyped by PCR using genomic DNA isolated from ear punches. Primer sequences and PCR conditions are available upon request.

### Behavioural Tests

#### BeamBreak

Spontaneous motor activity was tested in an open field test setup. Horizontal and vertical movement of mice was measured by infrared beams using home-made BeamBreak detector cages. Detector cages were cleaned with ethanol before experiments. Interruptions on the horizontal and vertical level were recorded for individual mice during a tracking period of 60 min. Interruptions on both levels accumulating over 60 min were normalized to control animals. All experiments were carried out between 1 and 5 pm.

#### Rotarod

Motor coordination was investigated by using the rotarod apparatus. The unit consisted of a rotating rod divided into five chambers by opaque synthetic walls. Power source and PC enabled speed and acceleration control of rotation. Soft tissue paper underneath the rod ensured a gentle landing after fall. Before each session, rod and walls were cleaned with ethanol, tissue papers were exchanged. Though the equipment had the capacity for five animals being investigated in parallel, motor coordination was detected in individual mice to exclude the impact of social stimuli on the animals’ performance. To solely test for motor coordination performance instead of motor learning, memory, or endurance, animals were not pre-trained and a short acceleration protocol was chosen^90^. Each mouse was transferred onto the rod with an initial speed of 5 rpm. At 10 s after placing the mouse onto the rod, the acceleration protocol was started (80 rpm final speed, 240 s ramp, acceleration rate of ∼20 rpm/min). Maximum speed and fall latency were noted. If an animal fell off the rod before starting the protocol, the actual time of fall was detected and another trial was allowed (up to three trials in total). In that case, the mean of all trials for time and speed was calculated instead of maximum values to correct for extra practice. If an animal fell of the rod within 5 s, poor placing by the experimenter was likely. Thus, another trial was allowed without counting the former one. All sessions were carried out between 2 and 5 pm.

#### Tail Suspension

The tail suspension test was used to test for depressive-like behaviour. The test was carried out according to^91^ with minor modifications. Before each session, mice were separately kept in a new cage for 30 min to acclimatize to the experimental room. Each mouse was investigated individually. Animals were suspended for a total time of 6 min by using laboratory tape (1.91 cm width, Roti®-Tape-marking tapes, Carl Roth). In order to prevent animals from climbing their tails, clear hollow home-made plastic cylinders were placed around their tails. Each session was recorded by a video camera. Active movement was defined as the condition when at least three paws were in motion. The time of immobility was calculated by subtracting the time of active movement from the total recorded time. Immobility time was normalized to the body weight at the time of the session and presented relative to control animals.

#### Sucrose Preference

K320E-Twinkle^DaN^ mice were inspected for anhedonic behaviour by using the sucrose preference test. Before the testing day, animals were transferred into a fresh open cage for 24 h together with their cage littermates. The open cage setup allowed offering two bottles at the same time, one with 5% sucrose solution and the other with tap water. After this habituation phase, littermates were transferred back to regular enclosed and individually ventilated cages. Experimental mice were kept single-housed in open cages for another 24 h. The amount of consumed water and sucrose solution was measured and presented in relation to the body weight at the time of the session. Values were normalized to control animals. To avoid influences by side-specific preferences, the bottles’ location was randomly changed (left or right). Mice were not deprived of food or water prior to the test. All experiments were conducted on weekends, ensuring a quiet environment for the tested animals.

#### Three Chamber Test

Sociability and preference for social novelty in K320E-Twinkle^DaN^ mice was investigated by using a home-made three chamber apparatus (19 L x 45 W x 23 H cm). The central chamber was connected to each neighbouring room by an open door, which could be closed with implementable and transparent plastic walls. Two mug-like metal containers (10 cm lower diameter, home-made) placed in the middle of each neighbouring room acted as separators between the exploring mouse (mouse to test) and the two unfamiliar animals (strangers). Thus, social interaction could be ensured while preventing physical violence. Experiments were recorded with a video camera. Within each experiment, animals of the same sex were used. Prior to each experiment, experimental mice and strangers were separately kept in a new cage for 30 min to acclimatize to test conditions. The whole apparatus, transparent walls, and containers were cleaned with ethanol before starting. Each experiment was divided into two sessions: the first to detect general social affiliation, the second to explore social novelty preference. The experiment was started with the exploring mouse placed in the central chamber and both doors blocked. After 5 min of habituation, the first session was started by removing both transparent walls. Hence, the exploring mouse had access to both neighbouring rooms with a container placed in the middle of each of them. Only one of the containers included a stranger. After 10 min of recording, the exploring mouse was transferred back into the central room and the doors were closed again. The second session started with additionally placing a new stranger under the second container. After another 10 min of recording the experiment was finished. For both sessions, the time spent in each room was measured.

### Positron emission tomography (PET)

Mice were between 19 and 21.5 months old at the time of PET imaging. For [^18^F]FDOPA measurements, mice received 15 mg/kg benserazide (Sigma-Aldrich, Steinheim, Germany) intraperitoneally 60 min before tracer injection to block peripheral decarboxylation of [^18^F]FDOPA. Tracer injection and measurement procedure was different for the three tracers used (Supplementary Table 1). Mice were anesthetized with isoflurane (5% for induction, 2% for maintenance) in O_2_/air (3:7), and for [^18^F]FDOPA and [^18^F]MNI1126 injection a catheter was inserted into the lateral tail vein. Following intravenous [^18^F]FDOPA injection, mice were transferred to their home cage for awake tracer uptake (30 min). [^18^F]FDG was injected intraperitoneally, and mice spent the tracer uptake period (40 min) awake in their home cage as well. For the subsequent scan, mice were anesthetized again and placed on an animal holder (Medres, Cologne, Germany). The awake uptake period after tracer injection prevents dampening of [^18^F]FDG uptake by anaesthetics^92^. Body temperature was maintained at 37°C by using a heating mat. Eyes were protected from drying with Bepanthen eye and nose ointment (Bayer, Leverkusen, Germany). [^18^F]MNI1126 was injected i.v. at the start of the scan without an awake uptake period.

For all tracers, a 30 min PET scan in list mode was conducted using a Focus 220 micro PET scanner (CTI-Siemens, Erlangen, Germany) with a resolution at the centre of field of view of 1.4 mm. This was followed by a 10 min transmission scan using a ^57^Co point source for attenuation correction. After the scan was finished, mice woke up in their home cage. Data were reconstructed as summed images. Full 3D rebinning was followed by an iterative OSEM3D/MAP reconstruction algorithm with attenuation and decay correction. The resulting voxel sizes were 0.47 mm x 0.47 mm x 0.80 mm. All further analysis was performed with the software VINCI 5.21 (Max Planck Institute for Metabolism Research, Cologne, Germany). Images were coregistered to an MRI template, and standardized uptake value ratios (SUVR_CB_) were calculated by dividing each image by the value extracted from a volume of interest (VOI) in the cerebellum. In addition, two striatal VOIs (left and right) were drawn voxel-by-voxel covering the whole striatum as seen in the MR image.

### Viral tracing

For viral labelling of remaining SN dopamine neurons, 20-month-old K320E/K320E^DaN^ and control mice (R26-K320E-Twinkle^+/+^; DAT^cre/+^) were used. For anterograde tracing, Cre-dependent AAV-pCAG-FLEX-EGFP-WPRE (1.8×10^13^ particles/ml; Addgene viral prep #51502-AAV1) was unilaterally injected into the SN of anesthetized animals at 3.0 mm posterior to Bregma, 4.2 mm deep from dura, and 1.6 mm lateral to midline^93^. AAV-pCAG-FLEX-EGFP-WPRE was a gift from Hongkui Zeng^94^. For retrograde tracing, Cre-dependent pAAV-FLEX-tdTomato (2.1×10^13^ particles/ml; Addgene viral prep #28306-AAVrg) was injected into the same hemisphere at 0.7 mm anterior to Bregma, 3.0 mm deep from dura, and 1.8 mm lateral to midline. pAAV-FLEX-tdTomato was a gift from Edward Boyden. For all intracerebral injections, mice were initially anesthetized with 4% isoflurane and fixed in a stereotaxic frame (#504926, WPI, Germany) at 2% isoflurane. Lubricating ointment was applied to the animals’ eyes to prevent them from drying (Bepanthen, Bayer, Germany). Body temperature was monitored by a rectal thermometer (RET-3, WPI, Germany) and kept constant by using a heating mat (MediHeat, V1200) and temperature control device (ATC2000-220, WPI, Germany). After reaching surgical tolerance, the incision site was shaved (HS 61 Contura, Wella, Germany) and cleaned with povidone iodine. Incision was made approx. 1.5 cm centrally along the skull bone and craniotomy was performed with a micro drill system (#503599, WPI, Germany) above the target regions. By using a micro-injector (#UMP3T 1, MPI, Germany), a target volume of 0.5 μl was applied at 100 nl/min for anterograde tracing. For adequate delivery of the virus, the cannula remained for another 5 min at the injection site. Subsequently, a target volume of 2.0 μl was applied at 200 nl/min for retrograde tracing. After removing the cannula, the wound was sutured and animals could recover in a heat box. Mice were subsequently provided with analgesics and carefully screened for signs of atypical behaviour and pain, according to European animal welfare guidelines. Three weeks after viral injection, mice were perfused and brains prepared for downstream histological analysis.

### dLight1.1

#### Stereotaxic surgeries

20-month-old mice were initially anesthetized using 5% isoflurane and placed on a stereotaxic setup (Kopf Instruments). During the surgery, mice were maintained under anaesthesia with 2% isoflurane and the animals’ reflexes were controlled regularly throughout the procedure. Octenisept (Schülke) was used to sterilize the skin and, prior to the incision, Anesderm (Pierre Fabre) was applied locally. A small hole was then drilled in the skull of the animals above the target site for viral delivery and fiber implantation.

400 nl of AAV5-CAG-dLight1.1 (a gift from Lin Tian, Addgene viral prep #111067-AAV5) was delivered into the dorsomedial striatum using glass pipettes at 0.7 mm anterior to Bregma, 3.4 mm deep from dura, and 1.9 mm lateral to midline. After virus delivery, an optical fiber (400 µm diameter, 0.48 NA, Doric Lenses) was implanted above the injection site and secured using dental acrylic (Super-Bond C&B, Sun Medical). Animals received tramadol via drinking water (1 mg/ml) for two days before and three days after surgery as well as buprenorphine (0.1 mg/kg, intraperitoneal) and meloxicam (5 mg/kg, subcutaneous) as post-operative care.

#### Data collection

Data were acquired using a RZ5P lock-digital processor controlled by Synapse software (Tucker-Davis Technologies, USA). The sampling rate used was 1.0173 kHz. For light excitation, a UV LED 405 nm and a blue LED 465 nm were sinusoidally modulated at 211 Hz and 566 Hz, respectively. The two lights were merged into a minicube (FMC4_AE(405)_E1(460-4909)_F1(500-550)_S) and directed into a fiberoptic rotatory joint (FRJ_1x1_PT-400/430-0.48_FCM-CM3, Doric Lenses) that was connected to the optical fiber attached to the animal. Emitted light from the dopamine sensor was collected and measured by a femtowatt photoreceiver (2151, Newport, USA). Synapse software performed real-time demodulation of the received raw fluorescence. Synapse software also allowed the synchronization of a video camera (HD Pro Webcam C920, Logitech; 10 frames per second) with the photometry recording.

#### Recording session

Fiber photometry recording started 4 weeks after surgery to allow complete recovery from surgery and optimal virus expression. Animals were acclimated to the experimental setup for one week by being attached to the patch cord inside the behavioural box for ∼15 min each day. The experimental setup consisted of a small transparent box (11.5 cm x 14 cm x 12 cm) placed inside a larger sound-attenuating box with opaque walls. Thus, animals were isolated from the experimenter’s view and surroundings during the recordings. In the subsequent week, animals underwent a recording session that started with a 10 min baseline period during which animals were allowed to explore the experimental box without any external interference. Following the baseline measurement, animals were exposed to different stimuli known to trigger dopamine release. These stimuli included a sudden opening of the experimental setup door; the delivery of a familiar 20 mg chocolate-flavoured sucrose pellet (F07290, Plexx) through a tube that ran from the outside of the larger box to avoid interference with the animal, and exposure to a fresh home cage.

#### Data analysis

A custom Matlab script was used to perform the fiber photometry analysis. To remove artefacts associated with the start of the system, the first 10 s of each recording were discarded. Raw data were downsampled to 50 Hz. To correct for signal fluctuations originating from motion artefacts, the 405 nm isosbestic reference was aligned to the 465 mm dLight signal using a least-squares linear fit, and was then used as the baseline to calculate the 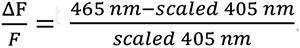 . For the analysis of peri-stimulus events, data were extracted using a time window centred around manually identified events following video analysis. For each type of stimulus, all trials from one animal were Z-scored and averaged. A time-window of -5 s to +10 s was used for the opening of the door and approaching of the pellets. The baseline used to calculate the mean and standard deviation was -5 s to -1 s. In case of exposure to a fresh cage, the time-window consisted of -60 s to + 300 s. Again, Z-score was calculated using the baseline -30/-60 s to – 1 s. All data were averaged across animals from the same group and the area under the curve during specified time windows was calculated to perform subsequent statistical analysis.

Of note, since photometry data are normalized to the baseline DA release within each subject, the set of results described above does not exclude the possibility of differences in basal, absolute, striatal dopamine levels. Consistent with the possibility of basal differences in dopamine release between the genotypes, the inspection of the raw demodulated dLight1.1 signals i.e., 465 nm signal, revealed striking disparities between the two groups (Extended Data Fig. 5g). All recorded K320E/K320E^DaN^ mice exhibited attenuated basal dopamine fluctuations as compared to control mice, pointing towards a reduced spiking amplitude. Importantly, these differences were not observed when analysing the reference 405 nm signal arguing that these variations do not arise from a technical artefact associated with changes in locomotion.

### Striatal tissue analysis

In order to obtain precise and pure striatal tissue for protein and RNA analyses, cryosections from 20-month-old K320E/K320E^DaN^ and control animals were used. Mice were sacrificed by cervical dislocation. Dissection equipment and embedding molds were treated with ethanol followed by RNaseZAP (Thermo Fisher Scientific). Freshly isolated brains were embedded in OCT compound (Tissue-Tek), fast-frozen on dry ice and stored at -80°C. 100 µm thick striatal sections from Bregma 1.18 mm to 0.26 mm were cut in coronal plane using a cryostat (Leica, CM3050 S). Sections were collected in an Eppendorf tube and kept chilled within the cryostat chamber at -20°C until the whole striatum was cut. To obtain rather pure striatal tissue and leave out frozen OCT compound, the striatum was framed by using a razor blade, releasing only the cut framed tissue. Collection tubes containing the whole striatum were stored at -80°C until further processing.

#### Western blot

Striatal tissue was lysed with RIPA buffer (150mM NaCl, 1% Triton-X1000, 0.1% SDS, 50mM Tris-HCl pH 8, 0.5% Na-deoxycholate) containing protease inhibitor (Roche). Protein concentration was measured by using the Bradford assay. After electrophoresis, proteins were transferred to a PVDF membrane previously activated with methanol. Membranes were blocked for 1 h (5% milk in TBS-0.1% Tween-20) and incubated overnight with primary antibodies. Following antibodies were used: polyclonal α-TH (1:200; Abcam, ab112), monoclonal α-DAT (1:1,000; Abcam, ab5990), monoclonal α-D1R (1:200; Santa Cruz, sc-33660), monoclonal α-D2R (1:200, Santa Cruz, sc-5303), monoclonal α-MAO-A (1:1,000; Abcam, ab126751), polyclonal α-MAO-B (1:500; Abcam, ab137778), monoclonal α-COMT (1:1,000; Abcam, ab126618), polyclonal α-GFAP (1:1,000; Merck Millipore, AB5804), polyclonal α-GAPDH (1:1,000; Merck Millipore, AB2302), and polyclonal α-Parkin (1:2,000; Abcam, ab77924). After washing (TBS-0.1% Tween-20), secondary goat anti-mouse, goat anti-rabbit, and goat anti-chicken HRP (1:10,000 each; Jackson Laboratory) were used according to primary antibodies. Primary α-DAT was detected by using secondary rabbit anti-rat (1;1,000; Dako, P0450). Upon visualization by using the ECL Advanced Chemiluminescence kit (GE Healthcare Life Sciences®, UK) according to the manufacturer’s protocol, images were visualized using a LAS500 CCD camera. For detection of D1R and D2R, higher secondary antibody concentrations were needed (1:5,000), followed by visualization using the SuperSignal West Pico PLUS Chemiluminescence kit (Thermo Fisher Scientific, USA).

#### Real-time PCR

RNA from striatal tissue was isolated using the RNeasy Mini Kit (Qiagen) according to the manufacturer’s instructions. Reverse transcription of extracted RNA was performed using the High Capacity cDNA RT Kit (Life Technologies). Designed primers used in this study are outlined in Supplemental Table 2. Amplification reactions (triplicates) were set up using Power up SYBR Green PCR Master Mix (Thermo Fisher Scientific) and qRT-PCR was validated with the QuantStudio 1 Real Time PCR system (Applied Biosystems, Thermo Fisher Scientific). Expression level of target genes was calculated using the comparative method of relative quantification (2^−ddCt^) and normalization to *Gapdh* amplification.

### Laser capture microscopy of dopaminergic midbrain neurons

#### Tissue sectioning and rapid TH staining

Brain isolation was conducted analogously to striatal tissue analysis. One day before laser dissection, thin coronal midbrain cryosections (12 µm) were cut at -20°C and transferred to RNaseZAP-treated PPS FrameSlides (Leica Microsystems, # 11600294). Six sections per FrameSlide, ranging from Bregma -2.92 mm to -3.40 mm in equal distance, were collected in vertical orientation. Sections were subsequently stored at -80°C.

Short TH immunohistochemistry immediately prior to laser dissection was performed according to^38^ with minor modifications. In this study, rabbit anti-TH primary (1:25; Abcam, ab112) and biotinylated anti-rabbit secondary antibody (1:25; Jackson ImmunoResearch, #711-065-152) were used.

#### Laser capture microscopy

TH-positive cells within the SN from Bregma -2.92 mm to -3.40 mm were captured using the Leica LMD7000 system. Cells were cut at 40x magnification while laser power was kept to a minimum. The laser was calibrated before each session. Detailed laser settings and cutting protocols are outlined in Supplementary Table 3-5. Cells fell into the cap of a sterile collection tube (0.2 ml PCR tubes, Eppendorf AG, Germany), containing 5 µl of collection buffer (NEBNext Single Cell Lysis Module, New England Biolabs). Noteworthy, shielding the Leica system with a cloak during laser dissection substantially improved cell output and prevented draft-induced deflections of single cells during drop. TH-positive cells were dissected from the SN only to gather adequate numbers of cells for RNA sequencing and to limit the procedure’s duration to a minimum with regard to potential RNA degradation. After cells were collected (∼275), another 5 µl of collection buffer were added to the cap, followed by pipetting up and down five to ten times. Eventually, lysates were spun down in a table centrifuge (Corning Labnet Spectrafuge C1202-230V)). Presence of laser-captured cells in the cap of the collection tube was monitored before and after spin down (Extended Data Fig. 7a-c). Tubes were carefully sealed with laboratory film (Parafilm), labelled, and snap frozen on dry ice. The whole procedure from thawing the sections until snap freezing of collected cells never took longer than 2 h 30 min. Samples were kept at -80°C for further processing.

### RNA sequencing of laser-captured SN dopamine neurons

#### RNA extraction

RNA from laser-captured dopaminergic SN neurons was extracted by using the Arcturus PicoPure RNA Isolation Kit (Thermo Fisher Scientific) according to the manufacturer’s protocol with minor modifications. 10 µl extraction buffer were added to the cap of the tube, followed by pipetting up and down ten times. Another 40 µl of extraction buffer were applied into the tube. After 1 min spin down, samples were incubated for 30 min at 42°C. Eventually, purified RNA was eluted in 12 µl elution buffer and stored at -80°C.

#### cDNA and sequencing library preparation

Isolated RNA was concentrated in a speedvac to 8 µl volume. Library preparation was performed with the NEBNext® Single Cell/Low Input RNA Library Prep Kit for sequencing, including cDNA synthesis (reverse transcription with polydT primers, template switching and cDNA amplification with 21 cycles), fragmentation, end repair, A tailing, adapter ligation, and library PCR with dual barcodes (8 cycles) and was followed by library quantification (Qubit, Tape Station) and equimolar pooling of the individual libraries. The library pool was then quantified using the Peqlab KAPA Library Quantification Kit and the Applied Biosystems 7900HT Sequence Detection System and sequenced on an Illumina NovaSeq6000 sequencing instrument with an PE100 sequencing protocol.

#### Read mapping and gene expression profiling

Adaptors were filtered out using Flexbar^95^, and reads were mapped to the mouse reference genome GRCm39 with the STAR alignment tool^96^. Reads were counted and assigned to genomic features using FeaturesCount^97^. Differential gene expression was calculated with EdgeR^98^. Genes were determined as differentially expressed with a false discovery rate cut-off of 0.05. Metascape (http://metascape.org)^99^ was used to perform gene ontology and network analysis as well as for generation of Figure 6d and Extended Data Figure 6f. In particular, enrichment analysis was carried out in custom mode with KEGG pathway as ontology source. Terms with a p-value <0.05, a minimum count of 3, and an enrichment factor >1.5 were grouped into clusters based on their membership similarities. Within a cluster, the most statistically significantly term was chosen to represent the cluster.

### mtDNA analysis of laser-captured SN dopamine neurons

#### DNA extraction

DNA from laser-captured dopaminergic SN neurons was extracted by using the QIAmp DNA Micro Kit (Qiagen) according to the manufacturer’s protocol with minor modifications. 15 µl lysis buffer were added to the cap of the tube, followed by pipetting up and down ten times and 1 min spin down. Eventually, purified DNA was eluted in 20 µl elution buffer and stored at -20°C.

#### Sequencing

The entire mtDNA was amplified from extracted DNA with two overlapping long-range PCR amplicons^100^, tagged and indexed applying Illumina Nextera XT reagents and sequenced at high depth with Illumina Miseq v3 chemistry. Each DNA sample was sequenced twice, including independent PCR amplification, library preparation and sequencing. Sequencing reads were aligned to the mouse mitochondrial genome (NCBI37/mm9 and NC_005089 with mouse NumtS sequences masked) using BWA-MEM (version 0.7.15-r1142-dirty). Duplications and deletions were called through MitoSAlt (version 1.1) using the following conditions: score_threshold = 80; evalue_threshold = 0.00001; split_length = 15; paired_distance = 1000; deletion_threshold_min = 30; deletion_threshold_max = 30000; breakthreshold = -2; cluster_threshold = 5; breakspan = 15; sizelimit = 10000; hplimit=0.01; flank = 15; split_distance_threshold = 543. Mitochondrial single nucleotide variants (mtSNVs) were called using VarScan (version 2.3.9) and filtered at a 0.5% heteroplasmy threshold. Intersected mtSNVs of both technical replicates were kept for the final annotation.

#### Copy number measurement

mtDNA copy number was determined by droplet digital PCR targeting the mitochondrial genes ND1 and ND4 and normalised by cell count^101^. Droplet analysis was performed using the BioRad QX manager software and the log2 fold change of copy number between groups was calculated.

### Histological analysis and immunostaining

#### Immunohistochemistry

Animals were intracardially perfused with PBS (GIBCO; 140 mM NaCl, 10 mM sodium phosphate, 2.68 mM KCl, pH 7.4) for 2 min followed by 4% PFA in PBS for 15 min. Upon post-fixation in 4% PFA for 2 days, brains were dehydrated in a series of increasing ethanol solutions (Leica ASP300, CMMC Tissue Embedding Facility, Cologne, Germany) and embedded in paraffin (Leica EG1150 H, CMMC Tissue Embedding Facility). Coronal 5 µm sections were cut using a microtome (Leica RM2125 RTS). For immunohistochemistry, perfused paraffin-embedded sections were deparaffinised in xylene, washed in a decreasing series of ethanol solutions and washed in ddH_2_0. For epitope retrieval, sections were heated in citrate buffer (10 mM citric acid monohydrate, pH 6) using a microwave oven. After cool down, sections were washed (TBS), quenched with 0.3% H_2_O_2_/TBS solution and washed again (TBS). Subsequently, sections were blocked (10% normal goat serum in TBS) and incubated with primary antibody in carrier solution (3% skim milk powder/TBS) at 4°C overnight. Polyclonal α-TH (1:750; Abcam, ab112) and monoclonal α-DAT (1:500; Abcam, ab5990) were used in this work. After washing (TBS), secondary biotinylated antibody in carrier solution was applied for 1 h (donkey anti-rabbit, 1:500; Jackson ImmunoResearch, #711-065-152; donkey anti-rat, 1:500; Jackson ImmunoResearch, #711-065-153). After incubation with avidin/biotin solution (Vectastain Elite ABC HRP Kit, Vector Laboratories) and another washing step, sections were incubated with DAB solution (DAB Kit, Vector Laboratories). Incubation times with DAB solution were kept similar for all intra-experimental groups. In order to prevent differences in DAB staining intensity between groups due to methodological reasons, section order for DAB incubation was randomized. Following a short washing step in ddH20, sections were dehydrated in a decreasing alcohol series, cleared in xylene and eventually mounted with Entellan (Merck Millipore). Upon curing of mounting medium, sections were imaged by using a slide scanner equipped with a 40x objective (SCN400, Leica, and S360, Hamamatsu).

#### Stereological quantification of dopaminergic neurons

Stereological quantification of TH-positive cells was conducted using the physical disector method^102, 103^. In this established approach, two adjacent, thin sections are used, in which the target particles are only counted if they appear in one of the sections. Furthermore, target particles are counted in randomized areas using a counting frame, in which they are only considered when touching the open frame, whereas those touching the forbidden frame are disregarded. In our case, target particles were classified as cells showing TH immunoreactivity with identifiable nucleus. TH-positive cells were counted within the allowed frames located in the SN and VTA. In total, nine pairs of adjacent midbrain sections covering Bregma -2.54 mm to -3.80 mm were chosen in equal distance. The number of counted TH-positive cells of all levels were summed and defined as dissector particle ‘Q’. Height sampling fraction ‘hsf’ 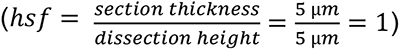 as well as area sampling fraction ‘asf’ 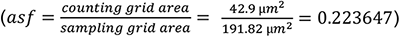 were kept constant. Section sampling fraction ‘ssf’ was set by the number of sections between each pair of sections (typically ∼20-30 sections, 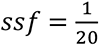, e.g.).

To ensure covering the same midbrain range despite potential differences in actual brain size between animals, the first and last slide was chosen uniformly for each animal followed by according calculation of the residual slides in between. The actual estimated number of TH-positive cells N was finally calculated by the formula: 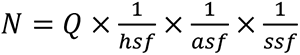.

#### Densitometric analysis of dopaminergic axon terminals

Striatal innervation of dopaminergic neurons was determined by optical density analysis using ImageJ. High resolution images were converted to RGB stacks and threshold was adjusted equally in all groups. Area fraction of TH- and DAT-positive fibres within the CPu and NAcc was analysed in two sections per animal at Bregma +0.74 mm. CPu and NAcc were defined according to the mouse brain atlas^93^. Optical density values from the corpus callosum were subtracted from matched striatal values to exclude unspecific TH and DAT signal.

#### COX-SDH staining

Visualization of Cytochrome c Oxidase (COX) deficiency was performed by enzymatic activity staining. COX is respiratory chain complex IV with subunits encoded by mtDNA, while succinate dehydrogenase (SDH), complex II, is entirely encoded by nuclear DNA. Impaired integrity of mtDNA results in COX-deficiency, while SDH-activity is sustained. Cells with decreased COX-activity stain blue for SDH, while cells with normal COX-activity appear brown.

Brain isolation was conducted analogously to striatal tissue analysis. Coronal midbrain cryosections (7 µm) were cut at -20°C and stored at -80°C until further processing. Sections at Bregma -3.08 mm were air dried and treated with COX incubation solution (100 μM Cytochrome C, 4 mM diaminobenzidine, 4400 U catalase in 0.1 M phosphate buffer) for 40 min at 37°C. After washing with ddH_2_0, sections were treated with SDH incubation solution (1.5 mM Nitroblue tetrazolium, 130 mM sodium succinate, 200 μM phenazine methosulfate, 1 mM sodium azide) for 180 min at 37°C. Sections were washed again, dehydrated using 95% and 100% ethanol, air dried and mounted with glycerol-gelatine. Sections were imaged by using a slide scanner equipped with a 40x objective (SCN400, Leica; S360, Hamamatsu).

#### Immunofluorescence in paraffin-embedded sections

Perfused paraffin-embedded microtome sections (5 µm) were generated analogously to immunohistochemistry. Upon deparaffinisation and epitope retrieval, sections were washed (TBS-0.5% Tween-20) and blocked with antibody diluent with background reducing components (Dako, Agilent). Primary antibodies in antibody diluent were applied overnight at 4°C. Following primary antibodies have been used: polyclonal α-TH (1:750; Abcam, ab112), monoclonal α-TH (1:500; Santa Cruz, sc-25269), monoclonal α-mtCOI (1:1,000; Abcam, ab14705), monoclonal α-NDUFB11 (1:500; Abcam, ab183716), monoclonal α-DAT (1:500; Abcam, ab5990), and polyclonal α-Bassoon (1:200, Synaptic Systems, #141 002). After washing (TBS-0.5% Tween-20), secondary antibodies in antibody diluent were applied for 2 h. Secondary goat anti-mouse and anti-rabbit were used according to primary antibodies (1:1,000; A11001, A32727, A21235, A32731, A21428, and A32733; Molecular Probes, Invitrogen). Secondary goat anti-rat (1:1,000; Alexa Fluor 647, Abcam, ab150167) was used for binding of primary rat α-DAT. Sections were washed (TBS-0.5% Tween-20), mounted (Fluoromount-G, Invitrogen) and dried at 4°C for at least one day before imaging.

#### Mitochondrial complex I and IV immunofluorescence

Fluorescence images of perfused paraffin-embedded midbrain sections at Bregma -3.08 mm were obtained using a Leica SP8 confocal microscope (Leica, Germany) with 40x oil objective. High resolution images of TH-positive cells from SN and VTA were taken from both hemispheres according to the mouse brain atlas^93^. Expression levels of mtCOI, the catalytic subunit of complex IV, were analysed using ImageJ. Somata of TH-positive cells except nuclei were defined as regions of interest. MtCOI mean gray value of somata was calculated and normalized to the nucleus of each cell, preventing fluorescent intensity deviations between and within sections due to methodological reasons. Data are presented as superplots, showing mean values for each animal in the front and single cell values in the background. Levels of NDUFB11, a subunit of mitochondrial complex I, were determined in an analogous way.

#### Active zone–like release sites in dopamine axon terminals

Super-resolution images of paraffin-embedded microtome sections following TH and bassoon immunoreactivity were acquired with a Zeiss LSM 880 microscope with an AiryScan detector (Carl Zeiss Microscopy), using a Zeiss 63X oil lens, NA 1.4, as the primary objective. Dual-colour excitation was performed with an argon laser for 488 nm and He 543 laser for 561 nm. The Airyscan detector was used in SR mode using all 32 pinholes, thus increasing the resolution to around 140 nm in x, y and z to capture the image. The image was reconstructed by pixel reassignment and deconvolution on the Zen Black platform (Carl Zeiss Microscopy).

The Final image was produced on ImageJ by background subtraction (rolling ball 50) and one run of the smooth filter. 3-5 regions of interest (ROIs) were imaged from the dorsal striatum in each slice with imaging volumes of ∼ 39 x 39 x 4 mm3. Quantification of bassoon clusters overlapping with TH-positive nerve terminals was performed in an automated manner with a macro script written for ImageJ. In brief, the script detects the regions positive for both bassoon and TH and uses the Analyse particle function in ImageJ to tabulate the number of colocalization spots. ROIs selected from the AiryScan images and flattened by maximum intensity projection (2 x 2 um) were used as inputs for the script. To enhance the degree of confidence, the bassoon channel was subjected to a 90-degree shuffling using a customized macro script on ImageJ. The overlap between shuffled images and unmodified images were hence compared and plotted using Prism.

Representative images of Z-stack sections acquired with a Zeiss LSM 880 microscope with an Airyscan detector (Carl Zeiss Microscopy) were created using Imaris (Oxford instruments, UK). In brief, the x, y, z section images were projected with a 45-degree perspective and a 3D grid to enhance the projection.

#### TH- and DAT-Colocalization

Fluorescence images of perfused paraffin-embedded midbrain sections at Bregma -2.80 mm, -3.08 mm, and -3.26 mm were obtained with a Leica SP8 confocal microscope (Leica, Germany) at 20x. TH- and DAT-expressing cells were quantified using ImageJ.

#### Immunofluorescence in perfused free-floating sections

After perfusion and two days of post-fixation with 4% PFA, brains were stored (PBS-0.05% sodium azide) at 4°C until further processing. One day before immunofluorescence, midbrain (50 µm) and striatal sections (100 µm) were cut using a vibratome (Leica VT1000 S, Leica Biosystems, Germany). Perfused free-floating vibratome sections were washed (PBS) and blocked for 2 h (10% NGS, 0.2% BSA, 0.5% Triton X-100 in PBS). All incubation steps were performed with shaking (∼130 rpm, KS 501 digital, IKA, Germany). After washing (PBS), sections were incubated with primary antibody in carrier solution (1% NGS, 0.2% BSA, 0.5% Triton X-100 in PBS) at 4°C overnight. The following primary antibodies have been used for free-floating vibratome sections: polyclonal α-TH (1:750; Abcam, ab112), monoclonal α-TH (1:500; Santa Cruz, sc-25269) and polyclonal α-GFP (1:1,000; Life Technologies, A-6455), and monoclonal α-TH (1:500; Santa Cruz, sc-25269). Sections were washed (0.2% Triton X-100 in PBS) and incubated with corresponding goat anti-rabbit/mouse secondary antibodies (1:1,000; A32731, A32733, and A21235; Molecular Probes, Invitrogen) for 3 h at room temperature. After washing (PBS), sections were transferred to objective slides (Superfrost Plus, Thermo Scientific) and mounted (Fluoromount-G, Invitrogen). Slides were stored in darkness at 4°C until imaging.

#### Analysis of vibratome sections

High resolution images of midbrain and striatal slices were acquired by a Leica SP8 confocal microscope (Leica Microsystems, Germany). Five sections were chosen for midbrain (Bregma -2.92 mm, -3.14 mm, -3.36 mm, -3.58 mm, and -3.80 mm) and striatum (Bregma 1.18 mm, 0.92 mm, 0.66 mm, 0.40 mm, and 0.14 mm) analysis. Using the Leica LAS X software, whole slice overview images were generated by merging single images acquired at 10x. Each of the single images was composed of z-stacks spanning a thickness of 8-20 µm. Maximum projections of whole slice overviews were used for analysis. Three z-stacks of 1024x1024 format were used for midbrain, five z-stacks of 512x512 format for striatum. Analysis of high resolution images was conducted using ImageJ. To confirm K320E-Twinkle expression in dopaminergic midbrain neurons by detection of downstream GFP, the number of GFP- and TH-positive cells was determined at 5 months of age. Viral tracing of late-age SN dopamine neurons was validated by quantification of EGFP- and TH-positive cells within the midbrain, and EGFP- and TH-positive fibres within the striatum, respectively. Specificity of viral labelling was determined by quantification of cells co-expressing TH and EGFP, and comparison with the non-injected hemisphere as intrinsic negative control. Injection accuracy was analysed by quantification of EGFP-expressing cells located in the SN compared to the VTA. In addition, densitometric analysis of EGFP- and TH-positive fibres within the CPu and NAcc was performed. Fibre density was determined analogously to immunohistochemistry of perfused paraffin-embedded sections.

#### Verification of fiber placement and virus injection (dLight1.1)

After perfusion and two days of post-fixation with 4% PFA, brains were cryoprotected in 20% sucrose in 0.1 M PBS. 30 µm striatal sections were cut by using a cryostat (Leica, Germany). Tissue was coverslipped with Vectashield mounting solution containing DAPI (Biozol VEC-H-1200). To visualize dLight1 fluorescence and fiber location, a Zeiss Imager M2 microscope and AxioVision 4.2 software (Carl Zeiss) was used. Images were acquired as tile scans of the whole section at 5x magnification.

### Data analysis

Data analysis was performed using GraphPad Prism 8 (GraphPad Software, Inc.). Quantified data are presented as mean + SEM. For all box plots used in this work, the centre line indicates the median, the mean is shown as “+”, the box borders present the first and third quartiles, and the whiskers indicate the data range. Relative data is shown as percentage of control animals. Values of sample size (n) refer to actual mouse numbers. Unpaired t test, one-way and two-way ANOVA with post hoc comparisons (Bonferroni post hoc test), respectively, were used to define differences between groups. The cumulative frequency of heteroplasmic mtSNVs between genotypes and ages was computed by two-sample Kolmogorov-Smirnov test. Significance level of 0.05 was accepted for all statistical tests.

Graphical illustrations were generated using Photoshop CS2 (Adobe Systems Software). Schematics (Fig. 1a, Fig. 4a, Fig. 5c, e, h, k, Fig. 6a, Fig. 7a and Extended Data Fig. 4a, h, j, l, n) were created with BioRender.com.

## Data availability statement

All data generated or analysed in this study are included in this published article and its supplementary information, or available from the authors upon request.

## Acknowledgements

We acknowledge Steffen Koch, Katrin Wollenweber, Robin Wolter and especially Katrin Lanz for their extensive help by preparing paraffin-embedded brain sections and genotyping of the used animal cohorts. We would further like to thank Prof. Olga Corti (Paris Brain Institute, Sorbonne University) for sharing the Parkin KO mouse, the Eva Hedlund lab (Stockholm University) for sharing protocols and experience on the quick TH stain, Prof. Matteo Bergami (CECAD, Cologne) for sharing his expertise on viral tracing, and Prof. Birgit Liss (Physiology, Ulm) and her team for extensive discussion and supply of the rotarod device. We wish to thank the Cologne Center for Genomics (CCG) for RNA sequencing, especially Janine Altmüller, and our mechanical engineers under former leadership of Harald Metzner and Hans-Josef Reimer for the in-house construction of the equipment used for behavioural assessments. For their continuous support concerning microscopy, we conclusively would like to acknowledge the CECAD Imaging Facility, especially Dr. Christian Jüngst.

## Author contribution

T.P. and R.J.W. designed the study, interpreted the data and wrote the manuscript. T.P. performed the experiments except otherwise indicated. K.M.R. contributed to tissue collection and data analysis of TH immunohistochemistry in the striatum. P.H. quantified midbrain dopamine neuron number in Parkin KO animals. Y.N. and P.C. performed mtDNA analysis.

H.E. and B.N. conducted PET scans. A.C. and S.S. performed dLight1.1 experiments. R.W.J., R.C. and J.P. carried out active-zone immunofluorescence. T.R. and B.B. performed read mapping and gene expression calculation of the sequenced transcriptome. Viral tracing experiments were supported by M.A. All authors commented or edited the paper.

## Ethics declaration

The authors declare no competing interests.

---

**Extended Data Figure 1.**
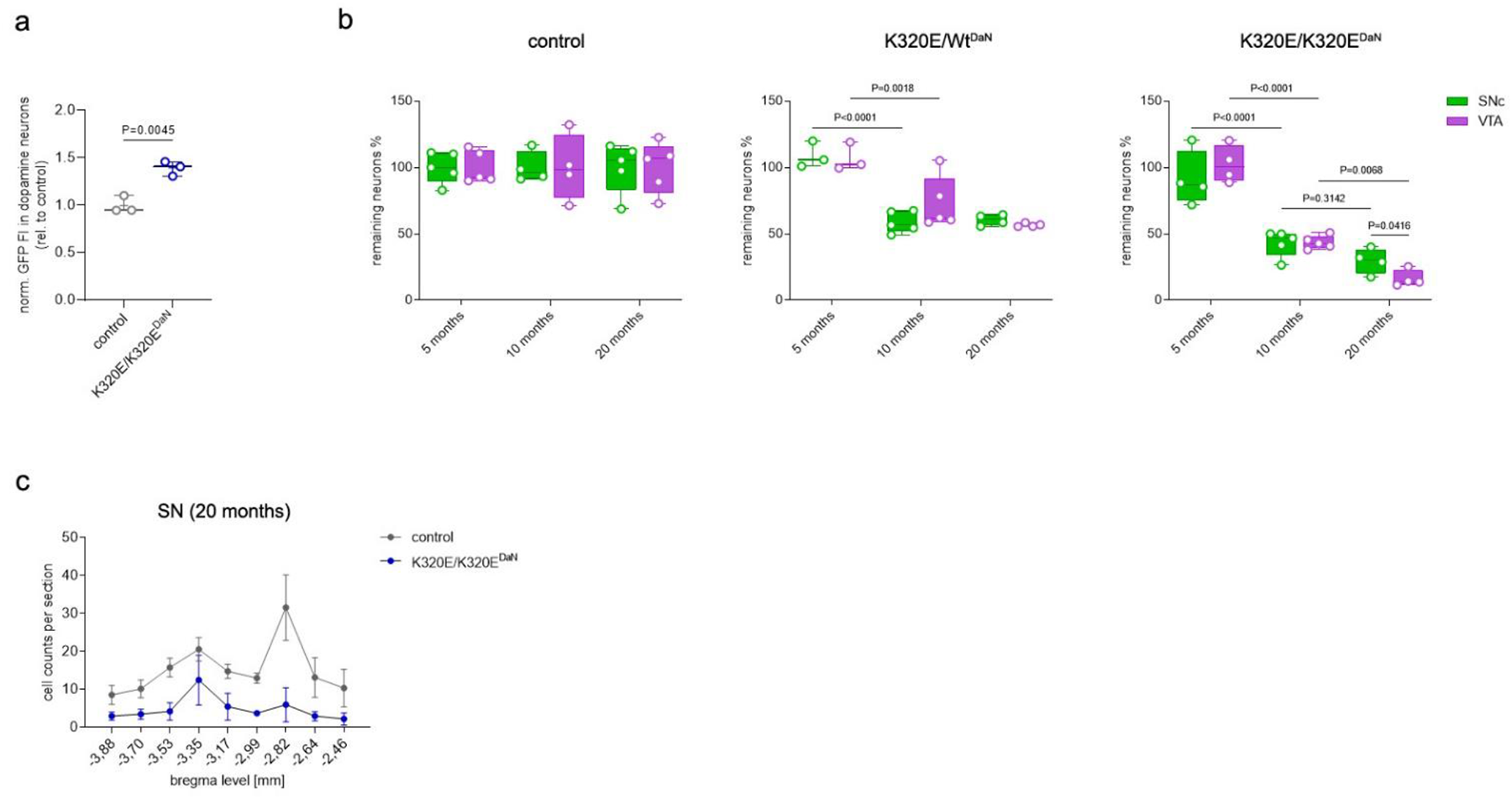
**a**, Normalized GFP fluorescent intensity of dopamine neurons from control and K320E/K320E^DaN^ mice at 5 months (n = 3). P value was calculated using unpaired two-tailed Student’s t test. **b**, Box plots showing the relative amount of remaining TH-positive neurons in age dependence (control, n = 4 5; K320E/Wt^DaN^, n = 3 5; K320E/K320E^DaN^, n = 4 5). **c**, Mean absolute counts of SN dopamine neurons in analysed sections from 20-month-old K320E/K320E^DaN^ and control mice (control, n = 5; K320E/K320EDaN, n = 4). P values were calculated using two-way ANOVA with Bonferroni’s correction for multiple comparison. Data are presented as mean ± s.e.m.

**Extended Data Figure 2.**
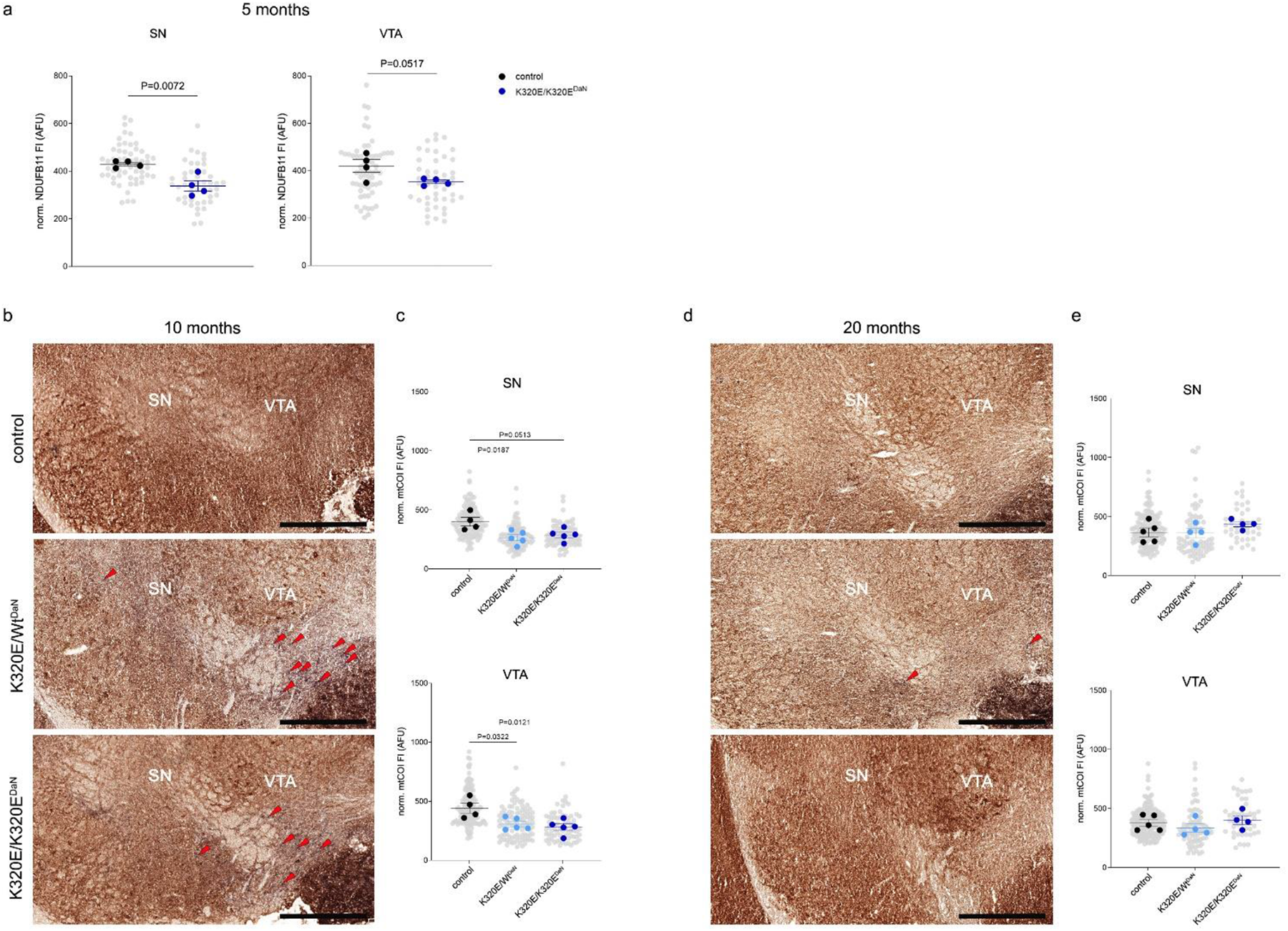
Normal expression and activtiy of cytochrome c oxidase in surviving dopamine neurons. **a**, Decreased normalized mean fluorescent intensity (FI) of NDUFB11 in SN and VTA dopamine neurons from K320E/K320E^DaN^ mice at 5 months (46 57 SN neurons, 44-60 VTA neurons, from 4 mice per group). P values were calculated using unpaired two-tailed Student’s t test. **b**, **d**, Representative images of COX/SDH enzyme activity staining in midbrain sections from control, K320E/Wt^DaN^, and K320E/K320E^DaN^ mice at 10 and 20 months; scale bar, 500 µm. **c**, **e**, Superplots presenting normalized mean fluorescent intensity of mtCOI in SN and VTA dopamine neurons at 10 (74-123 SN neurons, 73 139 VTA neurons, from 4 5 mice per group) and 20 months (39-145 SN neurons, 48 189 VTA neurons, from 4 5 mice per group). P values were calculated using one-way ANOVA with Bonferroni’s correction for multiple comparison. Data are presented as superplots including mean ± s.e.m.

**Extended Data Figure 4.**
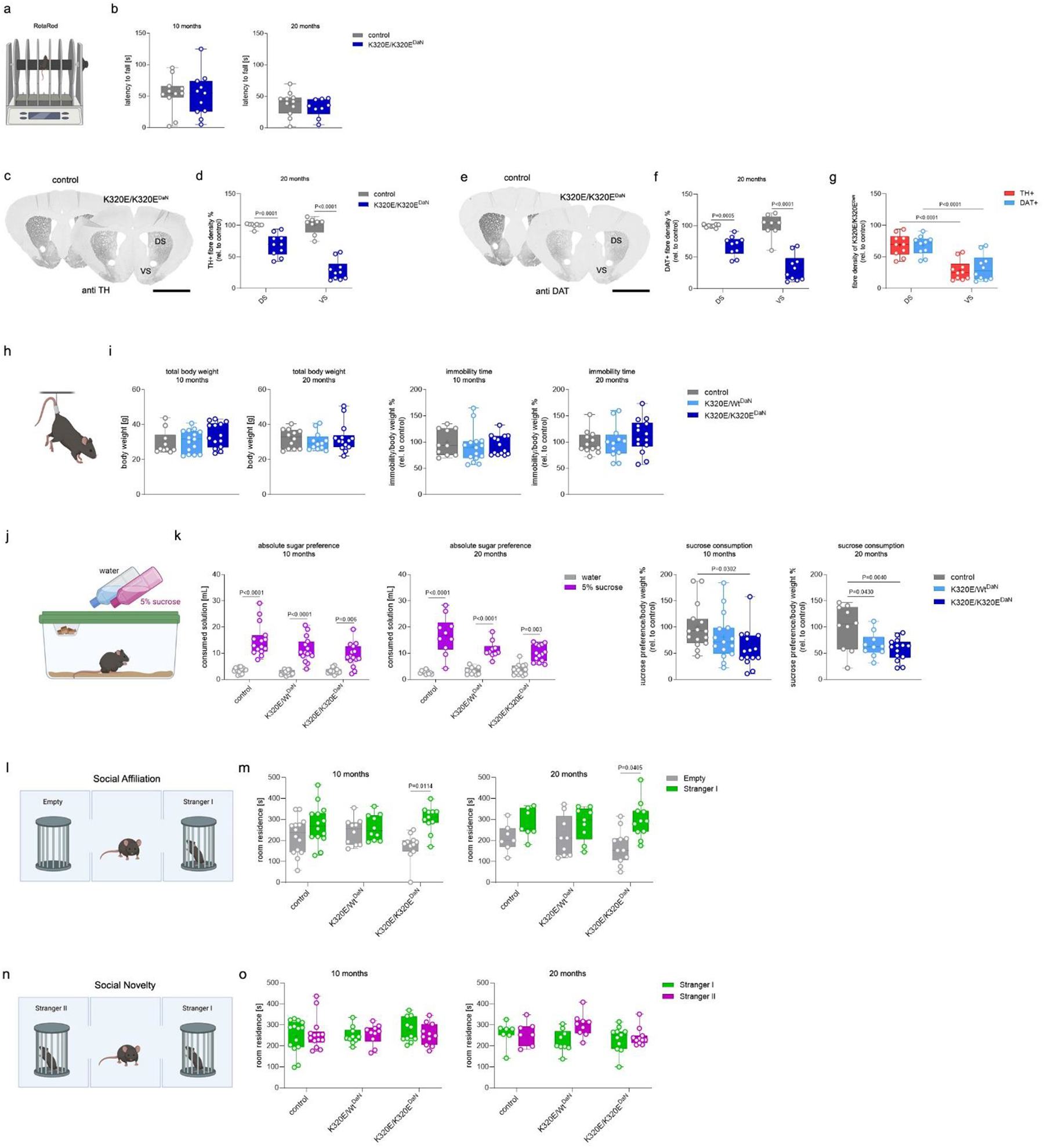
Denervation of the ventral striatum is associated with depressive-like and socially abnormal behavior. **a**, Schematic of the RotaRod setup. **b**, Latency time at fall from a rotating rod for control and K320E/K320E^DaN^ mice at 10 and 20 months (10 months, n = 11; 20 months, n = 9 11). **c**, **e**, Representative images of TH (**c**) and DAT (**e**) immunohistochemistry in the dorsal (DS) and ventral (VS) striatum of 20-month-old control and K320E/K320E^DaN^ mice; scale bar, 2 mm. **d**, **f**, Box plots presenting the quantitative analysis of TH-(**d**) and DAT-positive fibre density (**f**; n = 8 10). **g**, Comparison between the relative density of TH-and DAT-expressing fibres in the dorsal and ventral striatum of K320E/K320E^DaN^ mice. **h**, Schematic of the tail suspension test. **i**, Body weight and relative immobility time of control, K320E/Wt^DaN^, and K320E/K320E^DaN^ mice at 10 and 20 months (10 months, n = 10 15; 20 months, n = 12 14). **j**, Schematic of the sugar preference test. **k**, Absolute sugar preference and relative sucrose consumption at 10 and 20 months of age (10 months, n = 15; 20 months, n = 10 14). **l**, Schematic of the three chamber test measuring social affiliation. **m**, Box plots presenting the time each group spent in a room with an unfamiliar mouse placed under a container (stranger I) and in a room with an empty container, respectively (10 months, n = 10 14; 20 months, n = 7 11). **n**, Schematic of the three chamber test measuring social novelty. **o**, Box plots presenting the time each group spent in a room with another unfamiliar mouse placed under a container (stranger II) and in a room with the first unfamiliar mouse (stranger I), respectively (10 months, n = 10 14; 20 months, n = 7 11). P values were calculated using one- and two-way ANOVA with Bonferroni’s correction for multiple comparison Data are presented as mean ± s.e.m.

**Extended Data Figure 5.**
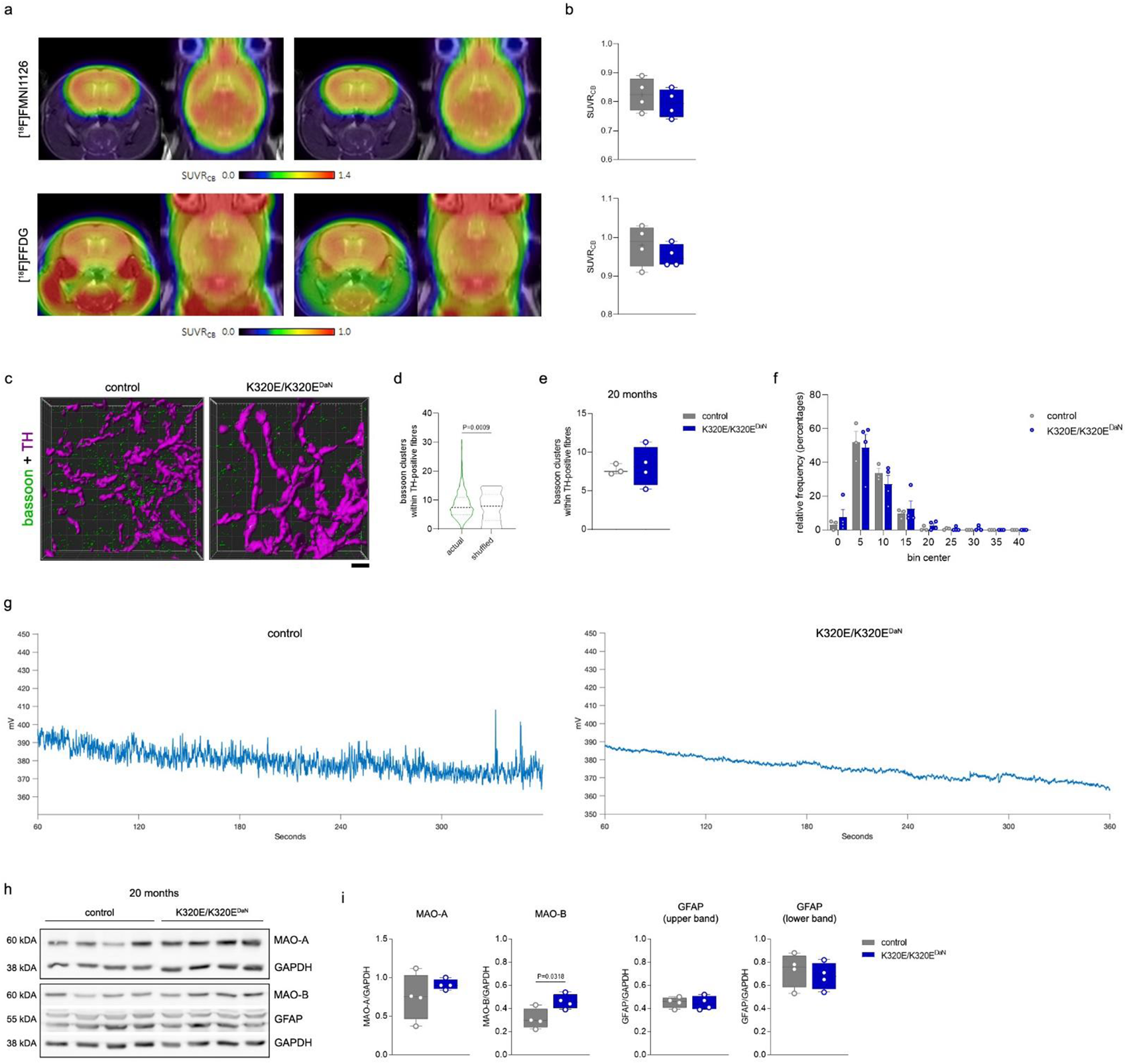
Compensatory striatal mechanisms. **a**, Averaged PET scan images presenting the uptake of synaptic density marker [18F]FMNI1126 and synaptic activity marker [18F]FDG by 20-month-old control and K320E/K320E^DaN^ mice. **b**, PET scan quantification showing the standardized uptake value ratios (SUVRCB) for [18F]FMNI1126 and [18F]FDG in the striatum. **c**, Representative 3D images showing the distribution of bassoon-positive clusters (green) and TH-positive axon terminals (magenta) in the dorsal striatum of 20-month-old control and K320E/K320E^DaN^ mice; scale bar, 1 µm. **d**, Quantification of all bassoon clusters within TH-positive fibres before and after shuffling of bassoon-positive objects (actual, n = 734; shuffled, n = 735). P value was calculated using unpaired two-tailed Student’s t test with Welch’s correction. **e**, Quantification of bassoon clusters within TH-positive fibres of control and K320E/K320E^DaN^ mice (control, 241 ROIs from 3 animals; K320E/K320E^DaN^, 493 ROIs from 4 animals). **f**, Relative frequency distribution of bassoon clusters within TH-labelled dopamine axons revealed no difference between control and K320E/K320E^DaN^ mice (control, n = 3; K320E/K320E^DaN^, n = 4). **g**, Representative traces of raw, demodulated dLight1.1 signals of control and K320E/K320E^DaN^ mice without external stimuli. **h**, **i**, Western blot analysis and quantification of indicated proteins in the striatum of control and K320E/K320E^DaN^ mice at 20 months. P values were calculated using unpaired two-tailed Student’s t test. Data are presented as mean ± s.e.m.

**Extended Data Figure 6.**
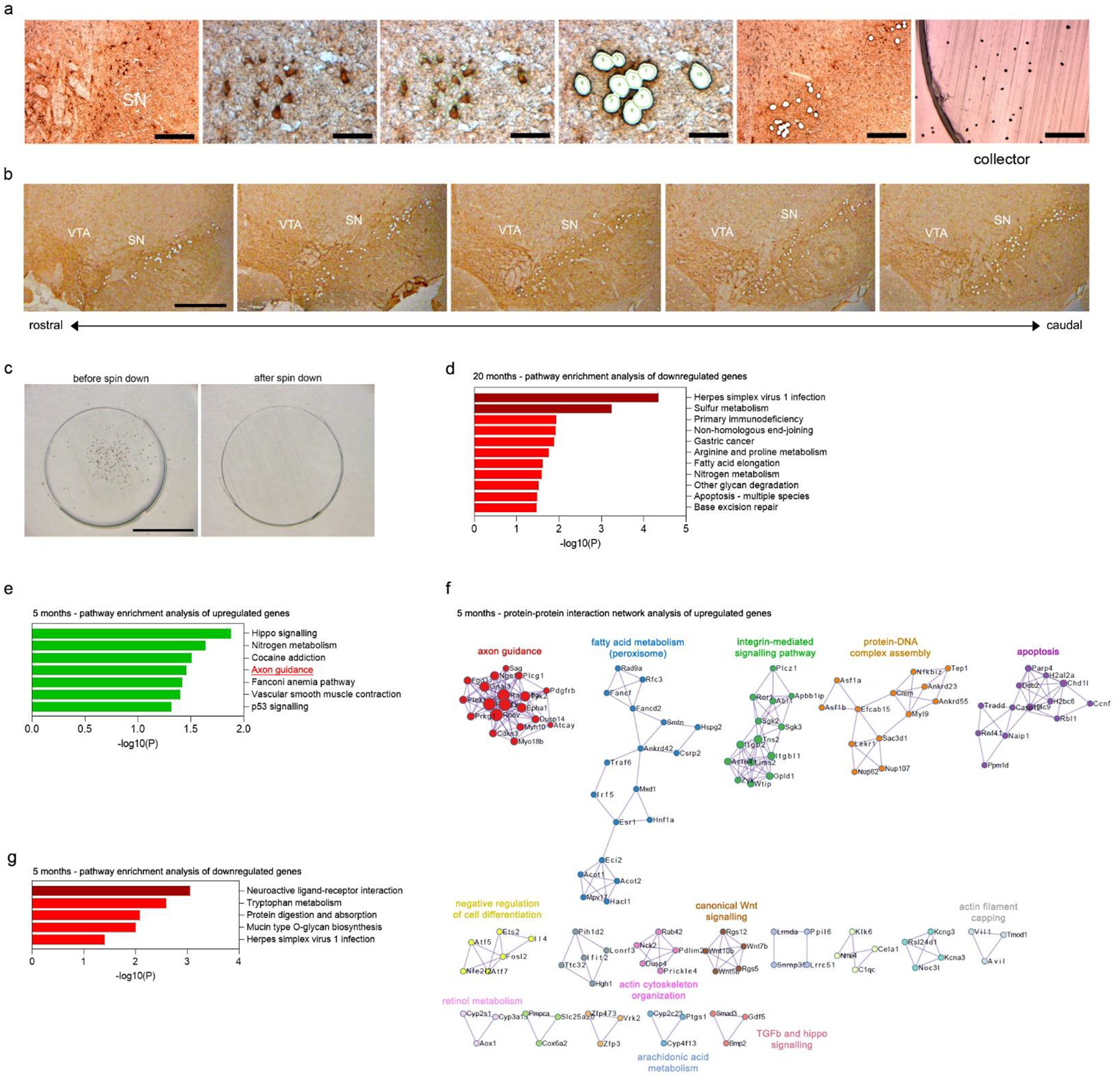
Laser capture microscopy of SN dopamine neurons and extended transcriptome data. **a**, Quickly TH-stained SN neurons from midbrain cryosections before and after laser dissection; scale bar SN overview before dissection, 200 µm; scale bar zoom, 50 µm; scale bar SN overview after dissection, 100 µm; scale bar collector, 100 µm. **b**, Representative immunohistochemical images showing laser capture microscopy of SN dopamine neurons along the rostro-caudal axis; scale bar, 500 µm. **c**, Cap-view before and after spin down confirms successful collection of SN dopamine neurons; scale bar, 1 mm. **d**, Bar graph illustrating enriched downregulated pathways in SN dopamine neurons of 20-month-old K320E/K320E^DaN^ mice. **e**, **g**, Bar graphs showing enriched upregulated (e) and downregulated (g) pathways in SN dopamine neurons of 5-month-old K320E/K320E^DaN^ mice. The most statistically significant clusters are shown; -log(10)P is the P value in negative log base 10. **f**, Network visualization of enriched protein-protein interaction for upregulated genes in 5-month-old K320E/K320E^DaN^ mice.

**Extended Data Figure 7.**
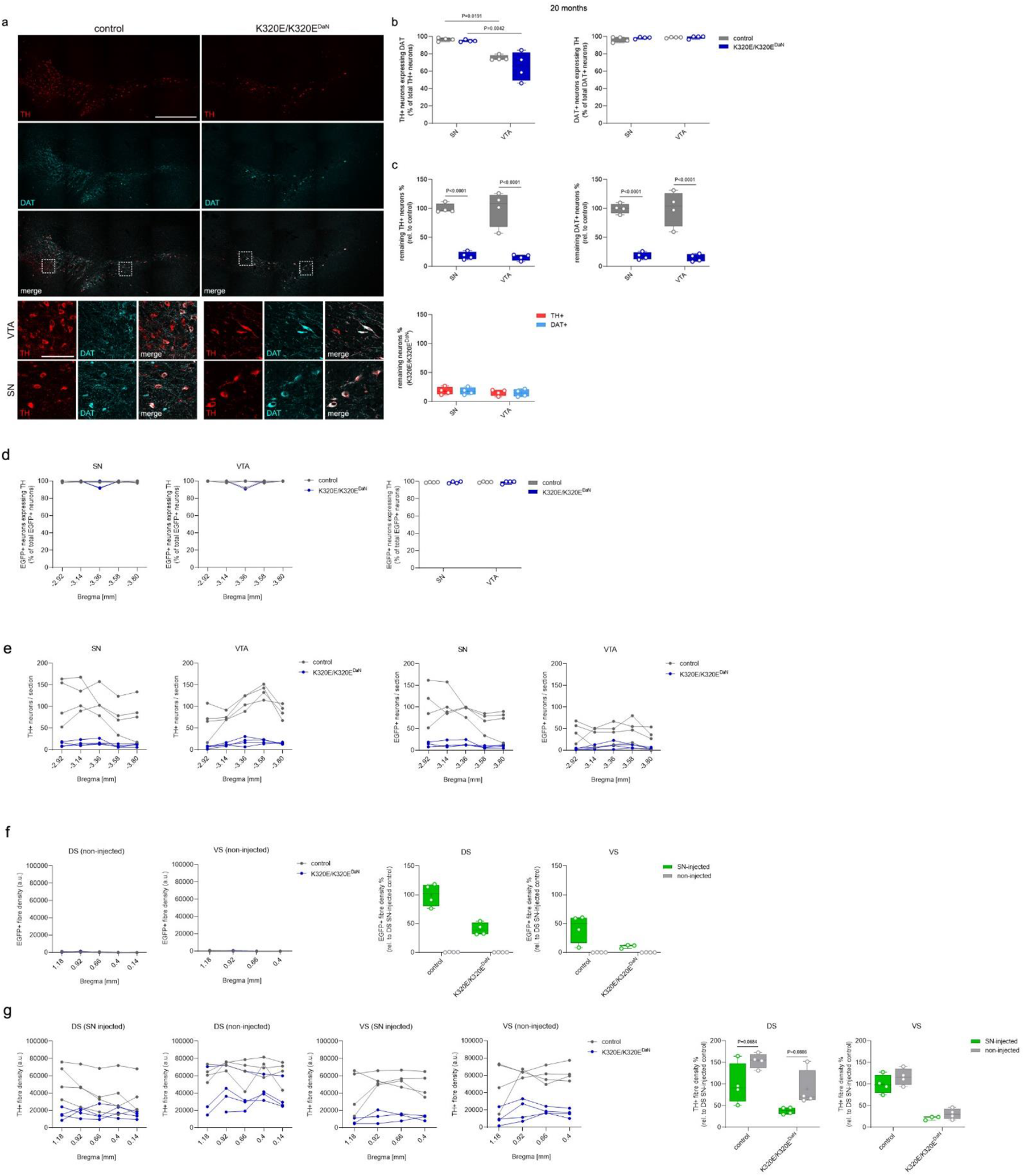
DAT expression in surviving SN dopamine neurons and extended viral tracing data. **a**, Representative immunofluorescent images revealing DAT expression in remaining TH-positive neurons of K320E/K320E^DaN^ mice at 20 months; scale bar, 500 µm; scale bar zoom, 100 µm. **b**, Quantification of TH-positive cells expressing DAT as well as DAT-positive cells expressing TH from control and K320E/K320E^DaN^ animals (3 distinct Bregma levels from 4 animals per group). **c**, Relative remaining TH-positive as well as DAT-positive neurons in the SN and VTA. **d**, Quantification of EGFP positive neurons expressing TH along the rostro-caudal axis upon viral injection into the SN (5 distinct Bregma levels from 4 animals per group). **e**, Total number of TH- and EGFP-positive neurons counted per Bregma level per group. **f**, Striatal EGFP-positive fibre density in the non-injected hemisphere and its comparison to the SN-injected side (DS: 5 distinct Bregma levels from 4 animals per group, VS: 4 distinct Bregma levels from 4 animals per group). **g**, Striatal TH-positive fibre density in the SN-injected and non-injected side. P values were calculated using two-way ANOVA with Bonferroni’s correction for multiple comparison. Data are presented as mean ± s.e.m.

**Extended Data Figure 8.**
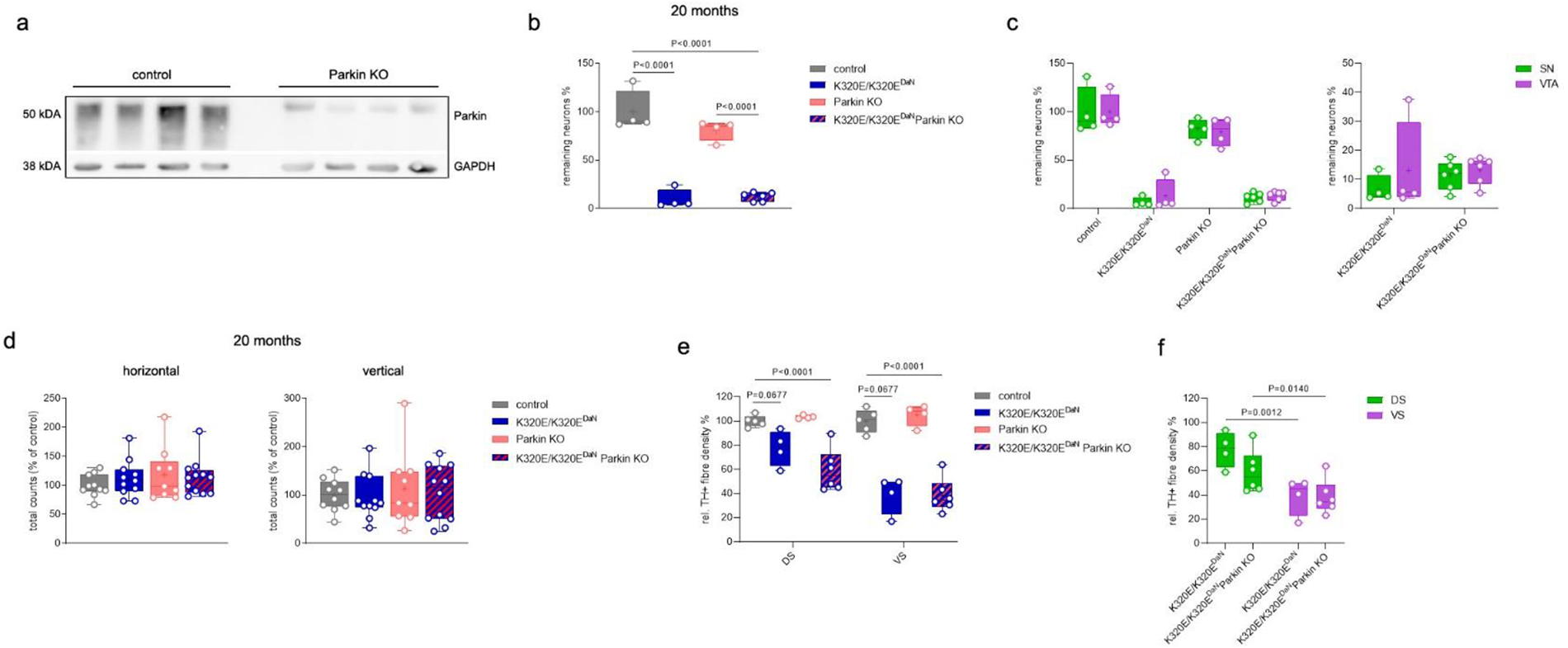
Loss of parkin does not worsen the phenotype of K320E-Twinkle^DaN^ mice. **a**, Western blot analysis confirming the lack of Parkin. **b**, Relative total TH-positive neuron number in K320E/K320E^DaN^ mice is not changed upon Parkin KO (control, n = 4; K320E/K320E^DaN^, n = 4; Parkin KO, n = 4; K320E/K320E^DaN^ Parkin KO, n = 6). **c**, No difference in remaining TH-positive neurons between SN and VTA. **d**, Similar total beam break counts for horizontal (locomotion) and vertical (rearing) movement of K320E/K320E^DaN^ and K320E/K320E^DaN^ Parkin KO mice over a period of 60 min (control, n = 10; K320E/K320E^DaN^, n = 11; Parkin KO, n = 9; K320E/K320E^DaN^ Parkin KO, n = 12). **e**, **f**, Parkin KO does not affect TH-positive fibre density in the striatum of K320E/K320E^DaN^ mice (control, n = 5; K320E/K320E^DaN^, n = 4; Parkin KO, n = 4; K320E/K320E^DaN^ Parkin KO, n = 6). P values were calculated using one- and two-way ANOVA with Bonferroni’s correction for multiple comparison. Data are presented as mean ± s.e.m.

